# CRISPR-mediated conditional mutagenesis of *Smad1/5/8* reveals BMP/GDF signaling restricts postnatal bone overgrowth

**DOI:** 10.64898/2026.01.22.701170

**Authors:** Rimma Levina, Alexander J. Weitzel, Alexander Y. Liu, Benjamin E. Low, Pierce W. Ford, Hayley Halaby, Hannah A. Grunwald, Erica G. Gacasan, Yasaman Moharrer, Robert L. Sah, Eric J. Bennett, Joel D. Boerckel, Michael V. Wiles, Kimberly L. Cooper

## Abstract

The BMP/GDF branch of TGF-β signaling regulates diverse aspects of skeletal biology, from skeletal development to maintenance and repair. However, the complexity, redundancy, and pleiotropy of BMP/GDF signaling have hamstrung a genetic dissection of its activities in different cell types over time. Here, we tested the feasibility of a three-transgene system using CRISPR/Cas9 to conditionally mutate six target sites, two each in the receptor-mediated *Smad1*, *Smad5*, and *Smad8* transcriptional effectors of BMP/GDF signaling. Briefly, we used *Prx1-*cre to activate a conditional Cas9 transgene by recombination in early limb bud mesenchyme; this endonuclease then complexes with gRNAs expressed from a polycistronic tRNA-gRNA array for targeted mutagenesis. Slower than expected accumulation of gRNA-directed mutations in each *Smad* produced an unexpected postnatal skeletal phenotype. Beginning around one month after birth, all animals developed hyperostosis on the surface of all long limb bones, which progressively worsened with age. This woven bone expansion occurred through proliferation of RUNX2+ osteoprogenitor cells in the cambium layer of the periosteum, producing an abundance of periosteal osteoblasts. Endosteal osteoblasts did not increase in number but increased their mineralizing activity. As a result, the marrow cavities narrowed, and the patella and carpal elements, which have no periosteum, increased internal bone mass without altering shape and size. Thus, while BMP/GDF signaling is known to promote early postnatal bone growth, these data support an additional homeostatic role during late postnatal osteogenesis by regulating both periosteal and endosteal osteoblasts. Although this genetically simple approach requires further optimization to improve efficiency, combining three transgenes produced more than 160 conditionally mutagenized animals with a fully penetrant and reproducible phenotype. This is an advance over traditional cre/lox systems that scale in complexity with the number of target loci, and it highlights the potential to model a wide range of genetically complex traits and disorders.

## Introduction

Each aspect of endochondral skeletal biology requires functions of the Bone Morphogenetic Protein (BMP) and Growth and Differentiation Factor (GDF) subfamily of the Transforming Growth Factor β (TGF-β) superfamily. However, despite the fact this signaling pathway has been a focus of intensive study for decades, partially overlapping redundancies and extensive pleiotropy have limited a genetic dissection of all its activities in different cell types and in different stages of skeletal development, maintenance, and repair from the embryo to aging adult^1^.

Dimers of fourteen BMP and GDF ligands bind to a heterotetrameric receptor complex comprising a dimer of one of four Type I receptors and a dimer of one of four Type II receptors^1^. The complexity of ligands and receptors converges to phosphorylate the receptor-mediated SMAD1/5/8 transcription factors, which drives their assembly and translocation to the nucleus, turning on a plethora of downstream target genes^1,2^. Consistent with the importance of these three transcription factors to transduce canonical BMP/GDF signaling, mice lacking *Smad1/5* or *Smad1/5/8* in *Collagen2*-cre+ chondrocytes develop with complete cartilage agenesis^3^. However, the severity of this phenotype and the combinatorial complexity of obtaining animals with a conditional triple knockout genotype, requiring seven alleles, precludes a deeper understanding of the varied roles of *Smad1/5/8* in other skeletal cell types.

Genetic complexity of pleiotropic gene networks is an obstacle to understanding a variety of biological processes from development to human disease. Here, to overcome the limitations of Mendelian inheritance, we developed a fully genome-encoded CRISPR/Cas9 transgenic system to conditionally mutate six target loci (two in each of *Smad1*, *Smad5*, and *Smad8*) using just three hemizygous transgenes. This work was inspired by our previous observation that the Rosa26^tm(CAG-LoxSTOPLox-Cas9,-eGFP)^ (Rosa26-LSL-Cas9^eGFP^) conditional transgene induced insertion and deletion (indel) mutations at 100% frequency in the male mouse germline when inherited together with the *Vasa*-cre transgene and a single *Tyrosinase* guide RNA (gRNA)^4^. The large number of available cre and chemically inducible cre transgenic mice would provide spatial and temporal resolution to any conditional CRISPR mutagenesis approach, as they have for recombination-mediated gene knockout. Lastly, a transgene expressing multiple gRNAs would complete a fully genome-encoded system to deploy conditional mutagenesis at any combination of multiple target loci.

Conditional Cas9 and Cas12a transgenic mice were initially used for delivery of pooled gRNAs by injection or viral transduction^5–7^. A recently reported conditional Cas12a system induced lung and pancreatic tumors after lentiviral transduction of gRNA arrays to mutate three tumor suppressor genes^8^. However, many tissues, including the heavily mineralized skeletal structures, are not amenable to injection and/or viral transduction. Instead, using a cre-regulated Cas and a genome-encoded gRNA array transgene would allow a simple breeding strategy to conditionally induce targeted mutations in inaccessible tissues and with cell type precision.

There are several possible strategies to multiplex gRNA expression. Each gRNA can be expressed using its own dedicated promoter, but multiple large promoters can make constructs unwieldy for cloning and transgene insertion. Alternatively, a single much smaller polycistronic transcript can allow individual gRNAs to be encoded into one construct and separated by a native CRISPR processing mechanism, as in the Cas12a system, or by a variety of encoded cleavage sites^9^.

In this study, we implemented a genome-encoded polycistronic tRNA-gRNA (PTG) array transgene to express six Cas9 gRNAs in mice. The PTG array is composed of alternating transfer RNAs (tRNAs) and gRNAs driven by a single RNA Polymerase III (Pol III) promoter. Endogenous RNA processing enzymes, RNaseP and RNaseZ, recognize tRNA secondary structure and cleave at specific 5’ and 3’ sites^10^. This mechanism of tRNA cleavage releases gRNAs from the PTG transcript. Additionally, each repeating tRNA contains internal promoter elements (box A and B) that recruit Pol III, potentially acting to enhance transcription^10^. Because each gRNA can be designed to target a different genomic sequence, a PTG array combined with a source of Cas9 expression can induce indel mutations at multiple target sites. PTG arrays have been delivered by transient transgenesis or transfection in several plant^10–12^, animal^13^, yeast^14^, and fungal^15–17^ species and in human^18,19^ and pig^20^ cell lines, but none have yet been encoded as a heritable transgene in mice. We predicted that the PTG array system could be adapted to efficiently generate conditional multi-locus knockout mice, because endogenous tRNA processing machinery is conserved across eukaryotes.

Here, we generated triple transgenic mice to mutate *Smad1/5/8* in limb bud derivatives. *Prx1-*cre initiates Cas9^eGFP^ expression by recombining the Rosa26-LSL-Cas9^eGFP^ transgene in early limb bud mesoderm, and a constitutively expressed PTG array encodes gRNAs targeting *Smad1/5/8*. We expected loss of BMP/GDF signaling during early limb development would cause total limb loss, so we were surprised when triple transgenic animals were born with normal limbs consistent with the near absence of indels at any target sequence at birth.

Instead, these mice gradually accumulated indel mutations postnatally and consequently developed extreme woven bone overgrowth (hyperostosis) along the surfaces of each long bone in all four limbs, which progressively worsened with age. Histology and immunofluorescence revealed that woven bone overgrowth initiates as an expansion of the cambium layer of the periosteum, including hyperproliferation of RUNX2+ osteoprogenitors that increase the number of osteoblasts. We also see increased mineralizing activity of endosteal osteoblasts. Consistent with these observations, the patella and carpals, which lack a periosteum, increase their internal bone mineral area without external woven bone overgrowth.

These data support dual roles of BMP/GDF signaling, suggesting the pathway promotes bone growth early and suppresses osteogenesis during later postnatal skeletal growth and maintenance. This could resolve seemingly contradictory phenotypes that resulted from ablation of different pathway components in different skeletal cell types and at different ages. For example, *Bmp2* ligand knockout in osteoblasts and osteocytes using *Osterix*-cre reduced cortical and cancellous bone volume^21^, but removal of *Acvr1* and *Bmpr1a* receptors individually in mature osteoblasts using *Collagen1a1*-cre increased bone mass^22–24^. Similarly, deletion of *Bmpr1a* in osteocytes using *Dentin matrix acid phosphoprotein 1* (*Dmp1-cre*) increased bone mass through elevated *Wnt* signaling and inhibition of RANKL and SOST^25^. The unique genetic manipulation, using moderately efficient conditional CRISPR mutagenesis, may preferentially reveal growth-suppressive functions of BMP/GDF signaling where loss of *Smad1/5/8* function gives mutant cells a competitive advantage in mosaic animals.

## Results

### Strategy to engineer tRNA-gRNA array for conditional mutagenesis

To select gRNAs for the PTG array, we designed four to target each *Smad* gene, initially referred to as Smad1_1-4_, for example (Fig. 1A). Each gRNA targets an exon encoding the conserved Mad Homology 1 (MH1) DNA-binding domain or the Mad Homology 2 (MH2) protein binding domain of each *Smad1/5/8* transcription factor. To test the mutagenesis efficiency at each target site *in vitro*, we collected GFP+ cells for genomic DNA extraction after transfection with a plasmid encoding Cas9, eGFP, and one of the twelve individual gRNAs. We then PCR amplified a ∼200 bp sequence around the gRNA target site, Sanger sequenced products, and analyzed results using the Inference of CRISPR Edits webtool (ICE, Synthego) (Fig. 1B)^26^. The ICE tool reports an overall indel frequency and the relative contributions of each unique indel sequence.

**Figure 1.**
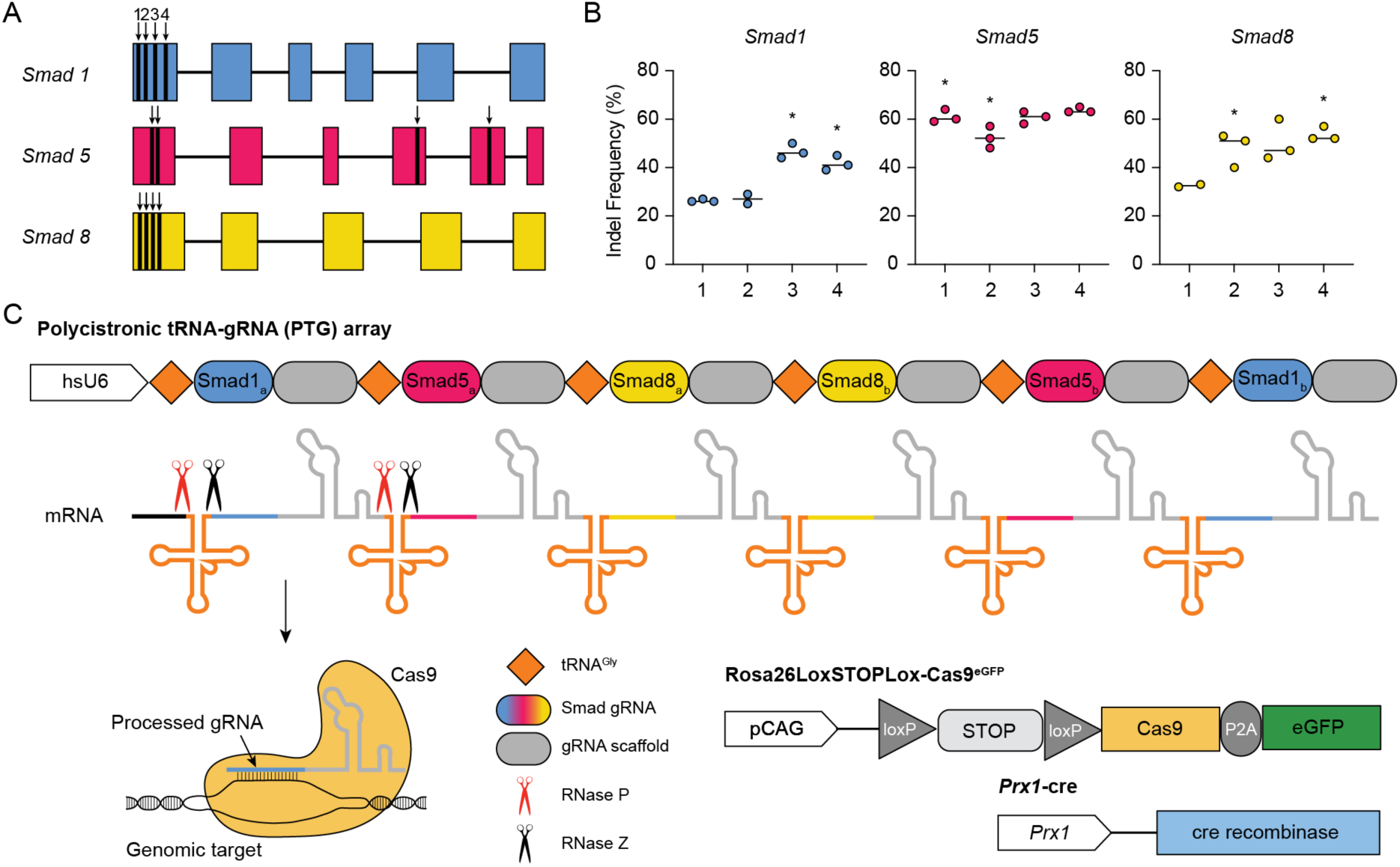
Three hemizygous transgenes were used to conditionally mutagenize Smad1/5/8. **(A)** Schematics of the exons in the *Smad1/5/8* genes, with black bars denoting each gRNA target site. Four gRNAs were designed per *Smad* gene targeting the MH1 domain (DNA-binding domain) or the MH2 domain (protein-binding domain). **(B)** Indel frequencies for each gRNA tested *in vitro* (n=3 each). gRNAs chosen for the polycistronic tRNA-gRNA (PTG) array are denoted with an asterisk (*). **(C)** Schematic of the three transgenes used for the breeding approach. The PTG array transgene, inserted into the Rosa26 safe harbor locus, is driven by the human hsU6 promoter. The PTG is composed of six total gRNAs interspersed with the rice glycine tRNA (tRNA^Gly^) sequence. Once transcribed, endogenous RNA processing enzymes RNase P and RNase Z cleave the tRNA sequence at specific sites, releasing individual gRNAs from the array. Each processed gRNA can then complex with Cas9 to induce double strand breaks. The Rosa26-LSL-Cas9^eGFP^ transgene, also inserted into the Rosa26 locus, is driven by a pCAG promoter. A P2A ‘self-cleaving’ peptide separates Cas9 from eGFP so that GFP can be used to track Cas9 expression. The conditional Rosa26-LSL-Cas9^eGFP^ transgene also encodes a floxed transcriptional STOP that is removed by the third transgene, *Prx1-*cre, for cell type conditionality.

Based on these data, we selected two gRNAs with the highest indel frequencies (>40%) for each *Smad* gene to assemble the PTG array (Fig. 1B, C). All selected gRNAs target the MH1 DNA binding domain, and none have a prevalent repair profile of three nucleotide insertion or deletion that would preserve the protein reading frame. Evidence suggests that multiple gRNAs with nearby targets increase the efficiency of CRISPR editing non-linearly, and either or both can induce loss-of-function mutations^27^. The two gRNAs targeting each *Smad* that are encoded in the array are referred to as Smad1_a_ and Smad1_b_, for example (Fig. 1C).

The rice glycine tRNA (tRNA^gly^) sequence has been used in rice^10^, human cells^18^, zebrafish^28^, and tobacco^11^, and its effectiveness was equivalent to species-specific tRNAs that were tested in zebrafish^28^, *Drosophila^13^*, and yeast^14^. We therefore assembled the PTG construct in an alternating tandem tRNA^gly^-gRNA (Fig.1C). Most prior studies have either not reported relative gRNA expression or activity from a PTG array or reported that the order of gRNAs in an array does not affect activity^28^. However, some studies have observed a correlation between position and mutagenesis, where gRNAs toward the 3’ end have lower indel frequencies than when tested individually^28,29^. Given this uncertainty, we encoded the gRNAs to begin with one for each *Smad* gene in the first three positions followed by the second gRNA for each in palindromic order (i.e., Smad1/5/8/8/5/1) to increase the chance that each target *Smad* gene would be mutated at least once. Conventionally, the U6 Pol III promoter is used to express gRNAs and was widely implemented in most PTG array construct designs^10,11,13,28,30^. Here, we used a human U6-Pol III (hsU6) to drive the PTG array.

We crossed PTG transgenic mice, inserted at the Rosa26 locus, with *Prx1-*cre transgenic mice to obtain double transgenic offspring (Fig. 2A). We then crossed Rosa26-PTG*; Prx1-*cre mice with homozygous Rosa26-LSL-Cas9^eGFP^ mice. Offspring that inherited all three transgenes (25% of each litter) will henceforth be referred to as PCC (Rosa26-<u>P</u>TG/ Rosa26-LSL-<u>C</u>as9^eGFP^; *Prx1-*<u>cre</u>) mice. Two-thirds of their littermates are control mice comprising two genotypes. PTG/Rosa26-LSL-Cas9^eGFP^ mice express *Smad1/5/8* gRNAs and inherited the conditional Cas9^eGFP^ transgene. These control for *Smad1/5/8* mutations that might arise due to promiscuous Cas9 expression in absence of cre. Rosa26-LSL-Cas9^eGFP^*; Prx1-*cre mice express *Cas9* in the *Prx1* lineage, but without gRNAs, and control for spurious gRNA-independent effects of Cas9 expression over many months. These also allow us to compare wild-type *Prx1-*cre lineage descendants with PCC lineage descendants, because eGFP is also expressed from the recombined Rosa26-LSL-Cas9^eGFP^ transgene. *Prx1-*cre mediates recombination in forelimb mesoderm at embryonic day 9.5 (E9.5) followed by hindlimb mesoderm at E10.5^31^. As expected, Cas9^eGFP^*;Prx1-*cre control mice express GFP strongly throughout derivatives of these cells including growth plate chondrocytes, inner and outer periosteal layers, endosteum, marrow cavity, embedded osteocytes, and articular cartilage^32^ (Fig. S1).

**Figure 2.**
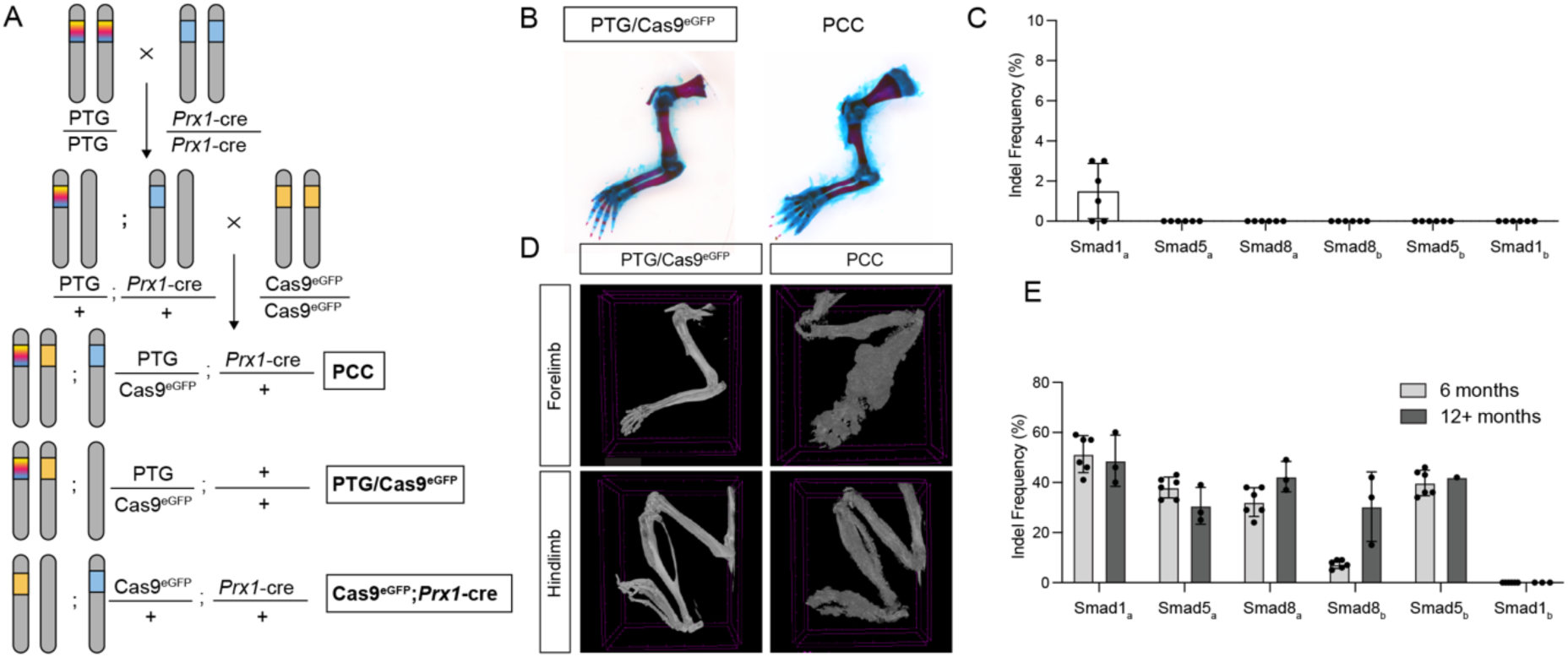
Mice that inherit all three transgenes exhibit postnatal hyperostosis. **(A)** Schematic of the breeding scheme used to unite all three transgenes (PCC) while also producing two classes of dual transgene littermate controls. **(B)** Alcian Blue/ Alizarin Red S-stained skeletal preparations of forelimbs from postnatal day 0 (P0) PCC and PTG/Cas9^eGFP^ animals (n=6 PCC, n=6 PTG/Cas9^eGFP^). **(C)** Indel analysis of bone tissue from the distal radius/ulna at P0 (n=6). **(D)** μCT scans of 12-month PCC and PTG/Cas9^eGFP^ forelimb and hindlimb. PCC limbs have substantial bone overgrowth. Purple dash marks in scans represent a scale of 1 mm in forelimbs and 2.5 mm in hindlimbs. All animals used for μCT at a range of ages show reproducible phenotypes: n=47 PCC, n=31 PTG/Cas9^eGFP^, n=29 Cas9^eGFP^;*Prx1-*cre. **(E)** Indel analysis of bone tissue from distal radius/ulna of six and 12+ month old animals (n=6 for six months, n=3 for 12+ months). Data represent as the mean +/-standard deviation.

Both conditional loss of *Smad1/5/8* in *Col2-*cre+ chondrocytes^3^ and *Prx1-*cre mediated deletion of the BMP receptors *Alk2*, *Alk3*, and *Alk6* cause complete limb agenesis^33^. We therefore predicted that efficient loss of *Smad1/5/8* in the *Prx1-*cre lineage would similarly produce mice without limbs and demonstrate clear proof-of-feasibility of the approach. However, triple transgenic PCC animals had no apparent skeletal deformities at birth (P0) (Fig. 2B). ICE analysis revealed <2% indel frequency at the first gRNA target site (*Smad1*_a_), and no indels were detected at sequential gRNA target sites (Fig. 2C).

We suspected that the extremely low mutagenesis could be caused by insufficient levels of gRNA expression from the single-copy genome-encoded transgene because there is evidence suggesting that indels accumulate over time, even in high-expressing *in vitro* systems. Once Cas9 induces a double strand break (DSB), it can be repaired perfectly or with an indel^34^.

Perfectly repaired sequences can be cut again, and thus repeat cutting accumulates the irreversible indel state over time^26,35^. Indel accumulation could be delayed even further by rate-limiting gRNA and/or Cas9 expression levels. Studies implementing similar PTG array strategies also observed a reduction in cutting efficiency from arrays containing a higher number of gRNAs when compared to arrays with fewer gRNAs, attributing it to the competition for Cas9 among several gRNAs at once^10,28^.

We therefore predicted that sustained gRNA and Cas9 expression over time in cells of the *Prx1-*cre lineage might allow mutations to accumulate in *Smad1*, *Smad5*, and *Smad8* producing a phenotype later during postnatal development. Indeed, by four months after birth, we began to see a phenotype in live animals; masses on the hands and feet were visible through the skin (not shown). By micro-computed tomography (μCT) at a range of ages, we identified these masses as an expansion of disorganized bone distributed along the full length of the forelimb and hindlimb of PCC mice, which progressively enlarged as the animals aged up to two years (Fig. 2D). Consistent with the gradually worsening phenotype, indel analysis of genomic DNA isolated from the distal radius/ulna showed indel frequencies at *Smad1/5/8* target sites approaching 60% over time (Fig. 2E). It is important to note that excised bone is composed of *Prx1-*cre lineage and non-lineage cells, but only cells from the *Prx1-*cre lineage express Cas9 and therefore could have mutations. That the last gRNA (*Smad1*_b_) never cuts is unexplained at this time; persistent low cutting efficiency of the *Smad8*_b_ gRNA, however, is likely caused by an unintended single base insertion during the array synthesis, a consequence of decisions made as the Covid-19 pandemic loomed in March 2020 (Supplemental Table 6).

We first evaluated the likelihood that the hyperostosis phenotype is caused by mutations in *Smad1/5/8*. Unfortunately, an antibody that detects the phosphorylated form of all three SMAD proteins does not reliably detect protein by immunostaining in ossified adult bone. Protein quantification using the same antibody in Western blots was highly variable (Fig. S2A), likely due to the mosaicism of induced mutations and differences in the cellularity of PCC bone compared with controls (see below). We therefore isolated mRNA from the distal radius and ulna of PCC and control mice for bulk RNA-Seq one month after birth when the phenotype is first detected. At this early phenotypic stage, we see significant downregulation of *Smad1* and *Smad8* mRNA expression in PCC bone, likely due to nonsense mediated decay, and Gene Ontology analysis shows significant enrichment of TGF-β and BMP pathway components among genes downregulated in PCC bone (Fig. S2B, C).

To detect off-target gRNA mutagenesis, we used three *in silico* prediction tools to identify genomic locations that might be recognized by the gRNAs in the array. There are three sites with two-base mismatches in our C57/B6 background (two for the Smad1_a_ gRNA, one for the Smad5_a_ gRNA), and no site has a single mismatch for any gRNA. We PCR amplified each of the two-base mismatch sites from distal radius/ulna genomic DNA of PCC animals after hyperostosis develops [3 months (n=6), 6 months (n=3), and 18 months of age (n=1)]. We did not detect any indel mutations at any of these most likely off-target sites in any sample (Supplemental Table 1). The phenotype only appears in mice that inherited all three transgenes, as both control genotypes develop normal limb skeletons (n=114 PTG/Cas9^eGFP^, n=69 Cas9^eGFP^*;Prx1-*cre). Although conditionally-expressed Cas9 in PCC mice independently induces mutations in cells throughout all four limbs, hyperostosis is fully penetrant and reproducible along the entire proximal-distal axis of each limb in 160 PCC animals. Altogether, it is therefore highly probable that the phenotype is caused by indel mutations in *Smad1/5/8*. One limitation of this approach, however, is a technical inability to determine the *Smad/1/5/8* genotype of each cell *in situ* to associate which gene(s) are mutated with their respective cellular phenotype.

### μCT quantification of hyperostosis in PCC limbs

To understand the earliest manifestation of bone overgrowth, we collected PCC and PTG/Cas9^eGFP^ mice at one, two, and three months after birth for μCT scanning and quantification. We focused on the radius because of its severe overgrowth in aged mice. At one month, PCC animals have an expanded trabecular network, disrupted cortical bone, and a narrower marrow cavity (Fig. 3A). This continues to worsen in two-month and three-month PCC animals, when the bone also expands outward by appositional growth along the length of the radius (Fig. 3B).

**Figure 3.**
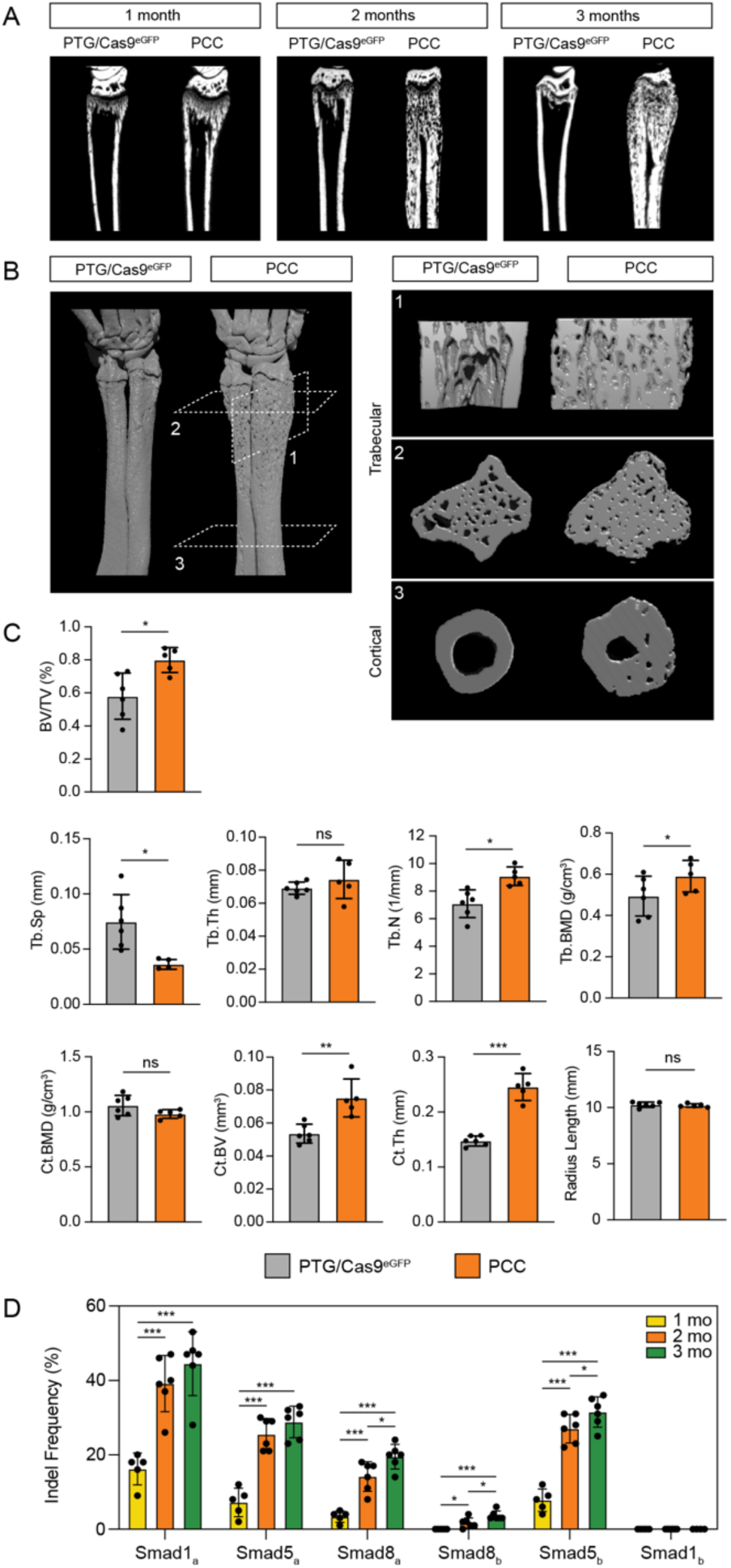
μCT analysis reveals that hyperostosis initiates by one month and progressively worsens. **(A)** Monthly time course μCT scans of the distal radius of triple knockout and control animals. **(B)** μCT reconstructions of the distal forelimb of two-month PCC and PTG/Cas9^eGFP^ mice. Top trabecular image corresponds to (1) – a transverse cross-section; bottom trabecular image corresponds to (2) – a longitudinal cross-section. Cortical bone transverse cross-section corresponds to (3). **(C)** Morphometric quantifications of cortical and trabecular regions of interest in **(D)** (n=6 PCC, n=6 PTG/Cas9^eGFP^). BV/TV = bone volume/total volume (%). Tb.Sp = trabecular separation (mm). Tb.Th = trabecular thickness (mm). Tb.N = trabecular number (1/mm). Tb.BMD = trabecular bone mineral density (g/cm^3^). Ct.BMD = cortical bone mineral density (g/cm^3^). Ct.BV = cortical bone volume (mm^3^). Ct.Th = cortical thickness (mm). Radius lengths of PCC and PTG/Cas9^eGFP^ mice are not significantly different (n=6 PCC, n=6 PTG/Cas9^eGFP^). **(D)** Indel analysis of bone tissue from distal radius/ulna of one-, two-, and three-month animals (n=5 one-month, n=6 two-month, n=6 three-month). Data represent mean +/-SD. * denotes Welch’s t-test <0.05. ** denotes Welch’s t-test <0.005. *** denotes Welch’s t-test <0.0005. ns = not significant.

Quantitative analyses of two-month PCC animals, compared with controls, showed significantly higher bone volume in both trabecular (BV/TV, %) and cortical bone (Ct.BV, mm^3^), as well as higher bone mineral density (BMD, g/cm^3^) in trabecular bone, but not in cortical bone (Fig. 3C). While there is a significant increase in trabecular number (Tb.N, 1/mm) and decrease in trabecular separation (Tb.Sp, mm), there is no significant difference in trabecular thickness (Tb.Th, mm) (Fig. 3C). We also observed a significant increase in cortical thickness (Ct.Th, mm) in PCC mice compared to controls (Fig. 3C). However, despite the importance of BMP signaling in growth plate maturation and bone formation^36,37^, PCC mice and controls have no significant difference in radius length (Fig. 3C).

To determine whether the earliest manifestation of expanded bone growth temporally coincides with accumulated mutations in *Smad1/5/8* we performed indel analysis of one, two, and three-month animals. At one month, the combined indel frequency for *Smad1* at both gRNA target sites was 16.2%, for *Smad5* it was 15%, and for *Smad8* it was 3.4% (Fig. 3D). This increased to 39.2%, 52.5%, and 15.8%, respectively, by two months and increased slightly again by three months. Together, these data suggest that *Smad1/5/8* mutations in cells of the *Prx1*-lineage cause an increase in trabecular and cortical bone mass.

### Cortical bone growth initiates as an expansion of the periosteum

To understand cortical bone thickening in PCC mice, we examined histology of the radius of one-, two-, and three-month-old animals at the distal and mid-diaphysis (Fig. 4). Cortical bone in both locations of one-month PCC mice appeared thicker when compared to PTG/Cas9^eGFP^ controls (Fig. 4A-B’’). At two-months, we also observed a thicker periosteum compared to controls (Fig. 4C-D’’). Typically, the periosteum is a thin membrane on the ossified surface of long bones. However, some regions of the periosteum extend into the bone of PCC mice. At three-months, pockets of cellularized tissue further disrupted cortical bone, and some pockets were embedded within mineralized bone (Fig. 4E-F’’). By contrast, histology at the endocortical surface appears similar to controls. There is no difference in growth plate morphology and heights of individual growth plate zones of PCC and control mice (Fig. S3).

**Figure 4.**
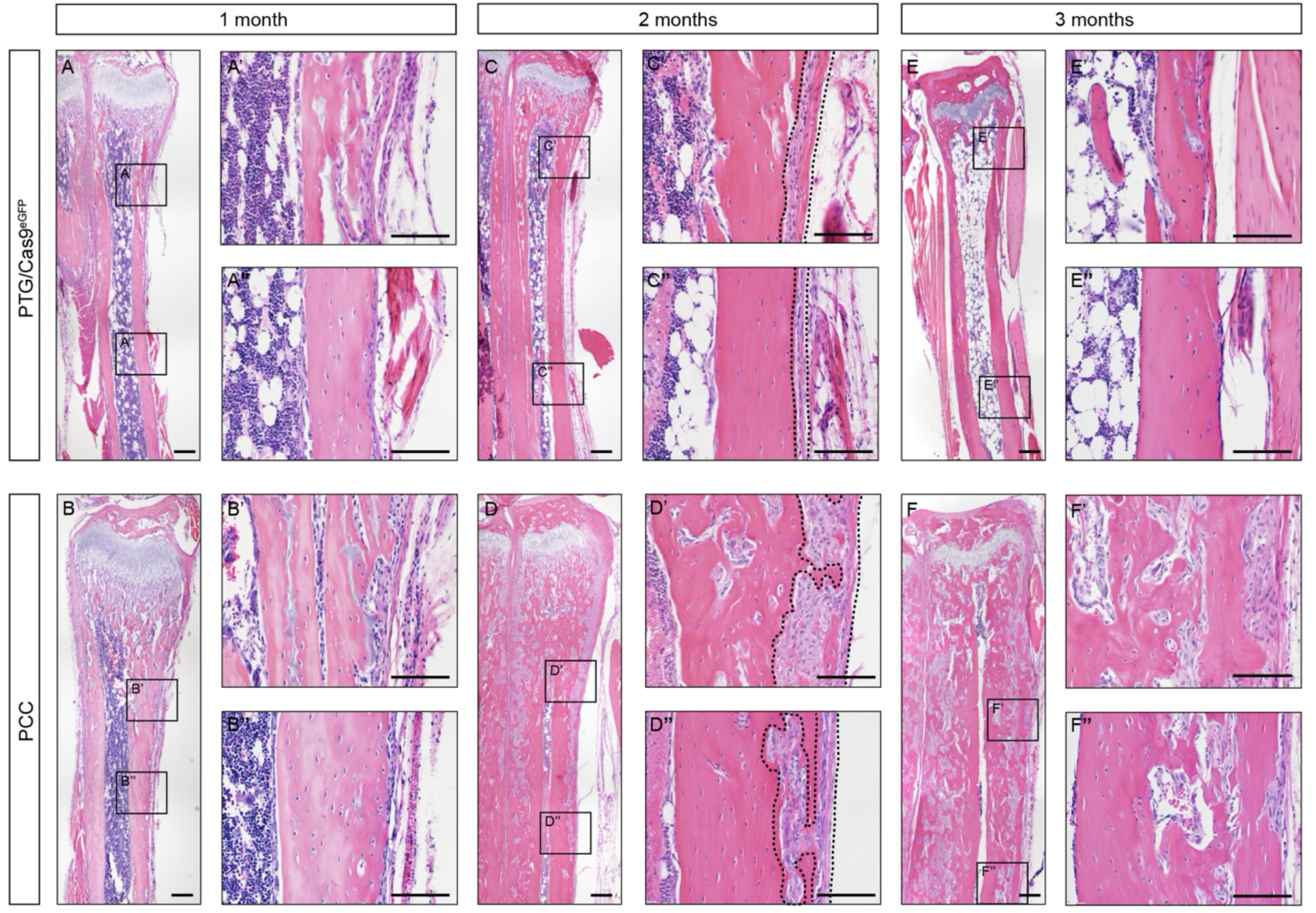
Histology time course reveals that woven bone overgrowth initiates with periosteal expansion. H&E staining of longitudinal sections. 4x overview of the distal portion of a **(A)** PTG/Cas9^eGFP^ radius and **(B)** PCC radius at one month. Boxed regions shown at a 20x magnification focusing on the **(A’, B’)** distal cortical bone and **(A’’, B”)** mid-cortical bone. **(C-C’’)** 4x overview and 20x magnifications of a PTG/Cas9^eGFP^ radius and **(D-D’’)** PCC radius at two months. **(E-E’’)** 4x and 20x magnifications of a PTG/Cas9^eGFP^ radius and **(F-F’’)** PCC radius at three months. Dashed lines in two-month panels outline the periosteum, highlighting expansion and disruption of compact cortical bone with disorganized periosteal tissue. Excess bone resembles woven bone. n=6 each for one-month PTG/Cas9^eGFP^ and PCC. n=6 each for two-month PTG/Cas9^eGFP^ and PCC. n=6 each for three-month PTG/Cas9^eGFP^ and PCC. 4x scale bars = 200 μm. 20x scale bars = 100 μm.

### Dynamic histomorphometry of bone mineralization in PCC mice

The disorganized bone in PCC animals resembles woven bone, a rapidly-deposited mineralized scaffold that is typically reorganized into lamellar bone^38^. Normal bone remodeling depends on an organized balance of bone formation and bone resorption, a harmony that is critical for maintaining correct bone mass, size, and shape. By contrast, PCC mice appear to accumulate woven bone appositionally without typical remodeling.

To determine how PCC mice have changed bone geometry, we performed dynamic histomorphometry by injecting animals at P37 with calcein (yellow) followed by Alizarin Red S (magenta) four days later and harvested tissue at P42 (six weeks postnatal; Fig. 5A). For these and subsequent analyses, we chose to focus on the mid-diaphysis of PCC and control animals. This region has a more simplified cortical structure compared to the re-organizing distal metaphysis, allowing us to better interpret the phenotype in cortical bone without the potential influence of expanding cancellous bone.

**Figure 5.**
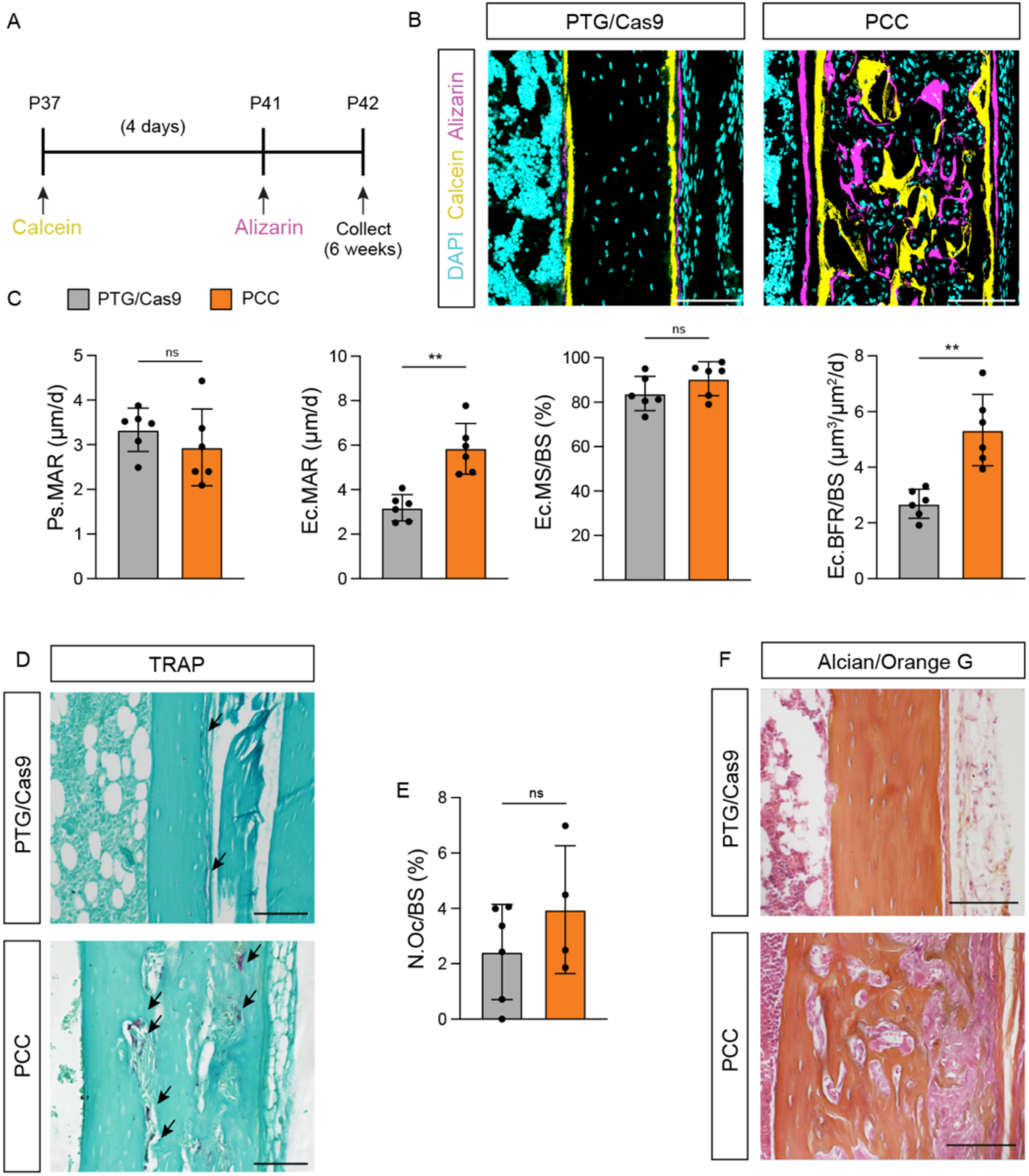
Dynamic histomorphometry of periosteal and endocortical bone growth. **(A)** Timeline of calcein and alizarin injections. Mice were injected with calcein at P37, then injected with alizarin four days later, and collected one day later at P42 (six weeks). **(B)** Representative images of dynamic histomorphometry at the mid-diaphysis in six-week-old mice (Calcein – yellow, Alizarin – magenta) (n=6 PTG/Cas9^eGFP^ and n=6 PCC). **(C)** Quantifications of mineralization, appositional growth, and bone formation. **(D)** Representative images at the mid-diaphysis for TRAP staining of longitudinal sections of the radius (n=6 PTG/Cas9^eGFP^ and n=4 PCC). Black arrowheads indicate osteoclasts. **(E)** Quantification of the number of osteoclasts normalized to bone surface. **(F)** Representative images at the mid-diaphysis for Alcian/Orange G staining of longitudinal radius sections (n=5 Cas9^eGFP^;*Prx1*-cre and n=6 PCC). Ps.MAR = periosteal mineral apposition rate (μm/day). Ec.MAR = endocortical mineral apposition rate (μm/day). Ec.MS/BS = endocortical mineralizing surface (%). Ec.BFR/BS = endocortical bone formation rate (μm^3^/μm^2^/day). N.Oc/BS = number of osteoclasts per bone surface. Scale bars (B) = 50 μm, (D) = 100 μm, (G) = 100 μm. Data represent mean +/-SD. * denotes Welch’s t-test <0.05. ** denotes Welch’s t-test <0.005. *** denotes Welch’s t-test <0.0005. ns = not significant.

We observed a massive increase in the combined periosteal/woven bone mineralizing surface in PCC mice compared to controls (Fig. 5B). However, the mineral apposition rate (MAR) at the periosteal surface was not significantly different (Fig. 5C). This suggests that osteoblast activity is normal, but there may be an increased number of osteoblasts contributing to excessive bone growth.

Some pockets of mineralizing surface within the cortical bone of PCC mice were lined with both labels while many contained only the second label (alizarin/magenta) (Fig. 5B). This could be explained by *de novo* labeling with alizarin or by resorption of mineral previously stained with calcein. We detected osteoclasts using tartrate-resistant acid phosphatase (TRAP) to evaluate bone resorption (Fig. 5D). There is no significant difference in the number of osteoclasts normalized to bone surface (N.Oc/BS) at the mid-diaphysis of PCC mice (Fig. 5E). However, we did observe osteoclasts residing within the cellularized pockets of PCC bone. Bone formation coupled with resorption that is mislocalized within the cortical wall may contribute to the overall disorganized woven appearance of bone.

To evaluate whether the disorganized bone involves a cartilaginous intermediate, as in the early fracture callus^32^, we stained bone sections with Alcian Blue/Orange G. We did not observe any chondrocytes marked with blue. Instead, staining primarily revealed bone and fibrous tissue (Fig. 5F).

Surprisingly, dynamic histomorphometry also revealed a significant increase in MAR and BFR/BS along the contiguous endocortical surface with no difference in MS/BS (Fig. 5C). This indicates an increase in osteoblast activity on the endocortical surface and suggests the endosteum also contributes to the observed increase in cortical bone thickness, though perhaps through a different cellular mechanism.

### Increased proliferation of RUNX2+ periosteal osteoprogenitors increases osteoblast number

Given the appearance of woven bone coincides with cellularized pockets with the histological appearance of periosteum, we next sought to confirm this tissue identity. The periosteum consists of two distinct layers - an outer fibrous layer composed of collagen fibers, fibroblasts, tendon attachments, and vasculature, and an inner cambium layer composed of mesenchymal and osteogenic progenitors and endothelial pericytes^39^. To detect the fibrous layer, we immunostained bone sections for Stem Cell Antigen-1 (SCA-1). We observed no difference in expression of SCA-1 in the fibrous layer of PCC mice compared to Cas9^eGFP^;*Prx1-*cre controls and no difference in fibrous layer thickness (Fig. 6A, B). To detect the cambium layer, we stained bone sections with Periostin (POSTN) and observed a marked increase in POSTN expression throughout the expanded tissue on the periosteal surface and within entrapped pockets (Fig. 6C). Together, this demonstrates an expansion of the inner cambium layer, and not the outer fibrous layer, of the periosteum in PCC mice.

**Figure 6.**
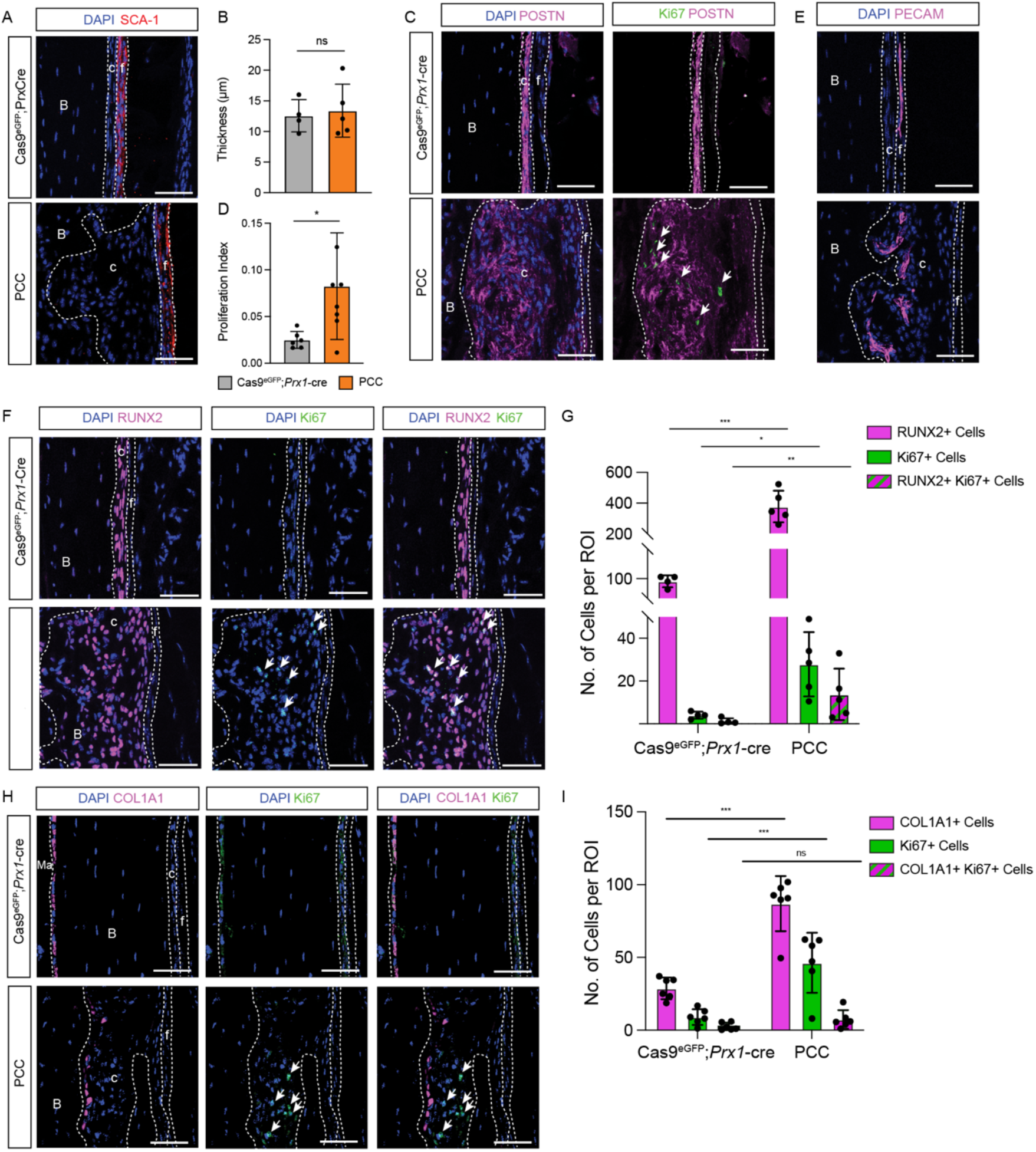
Inner cambium layer of the periosteum in PCC mice expands due to increased proliferation of RUNX2+ osteoprogenitors. **(A)** Representative images of SCA-1 immunostaining of longitudinal radius sections at two months focused on the mid-diaphysis of Cas9^eGFP^;*Prx1*-cre control and PCC mice (n=4 Cas9^eGFP^;*Prx1*-cre and n=6 PCC). **(B)** Quantification of fibrous layer thickness (μm) at the mid-diaphysis, ROI represented in (A). **(C)** Representative images for Periostin (POSTN, magenta) and Ki67 (proliferation, green) immunostaining of longitudinal radius sections at two months (n=6 Cas9^eGFP^;*Prx1*-cre and n=8 PCC). **(D)** Quantification of proliferation index (% Ki67+) at the mid-diaphysis, ROIs represented in (C). Significant periosteal expansion results from increased proliferation of cells in the cambium layer. **(E)** PECAM staining of longitudinal radius sections at two months (n=4 Cas9^eGFP^*;Prx1-*cre and n=5 PCC). **(F)** Representative images for RUNX2 (magenta) and Ki67 (green) immunostaining at the mid-diaphysis (n=4 Cas9^eGFP^;*Prx1*-cre and n=4 PCC). **(G)** Quantification of RUNX2+ (magenta bars), Ki67+ (green bars), and RUNX2+Ki67+ double positive cells (magenta and green striped bars) in ROIs represented in (F). **(H)** Representative images for COL1A1 (magenta) and Ki67 (green) immunostaining at the mid-diaphysis (n=6 Cas9^eGFP^;*Prx1*-cre and n=6 PCC). **(I)** Quantification of COL1A1+ (magenta bars), Ki67+ (green bars), and Col1a1+Ki67+ double positive cells (magenta and green striped bars) in ROIs represented in (H). B = bone. c = cambium layer. f = fibrous layer. Ma = marrow. Scale bars = 50 μm. Arrowheads indicate Ki67+ cells. Data represent mean +/-SD. * denotes Welch’s t-test <0.05. ** denotes Welch’s t-test <0.005. *** denotes Welch’s t-test <0.0005. ns = not significant.

To explore the cellular basis for the expanded cambium layer in PCC mice, we quantified cell proliferation and apoptosis. TUNEL staining detected a low frequency of cell death in both control and PCC mice (Fig. S4A). Kiel 67 (Ki67) immunofluorescence detected little to no proliferation in either the cambium or fibrous layers in control mice (Fig. 6C, D). However, PCC mice have markedly increased cell proliferation in the expanded cambium layer (Fig. 6C, D). To determine if Ki67+ proliferating cells in the periosteum of PCC mice express Cas9, and therefore could have *Smad1/5/8* mutations, we detected eGFP expression from the polycistronic Rosa26-LSL-Cas9^eGFP^ transgene. All Ki67+ cells are also GFP+ (Fig. S4B).

We next asked which population(s) of resident cambium layer cells are expanded and more highly proliferative. PRX1 is broadly expressed throughout the expanded cambium in *PCC* mice, suggesting this mesenchymal population might contribute to an increase in osteoprogenitors (Fig. S4C). Indeed, we also observed a significant increase in Runt-related transcription factor 2 (RUNX2) expressing osteoblast precursors in the cambium layer of *PCC* mice compared to controls (Fig. 6F). There is a 7-fold increase in the number of proliferating cells and a 4-fold increase in the number of RUNX2+ cells. Of all proliferating cells, 20% are RUNX2+ in control mice, which is increased to 56% in PCC mice (Fig. 6G).

Proliferative osteoprogenitors continually produce new osteoblasts by differentiation^40^. To determine if the highly proliferative osteoblast precursors contribute to an increased osteoblast number in PCC mice, we stained for Collagen1a1 (COL1A1). We observed a significant increase in the number of osteoblasts within the expanded cambium of PCC mice, as well as lining cellularized pockets embedded in cortical bone (Fig. 6H, S4D). Co-staining with Ki67 showed that the majority of COL1A1+ osteoblasts are not proliferative, consistent with their postmitotic state in controls; the few that are Ki67+ may have recently differentiated from a dividing pre-osteoblast (Fig. 6I). These data suggest that BMP/GDF signaling through *Smad1/5/8* suppresses proliferation of RUNX2+ osteoblast progenitor cells in the cambium layer of the periosteum. We hypothesize that pockets of cambium layer periosteum likely became ‘trapped’ within cortical bone, because they contain many osteoblast precursors that mature into osteoblasts and continuously deposit new bone, consistent with dynamic histomorphometry.

Additionally, both the fibrous and cambium layers are vascularized, though to different extents, and histology revealed blood cells within the entrapped cambium in *PCC* mice. We therefore stained for platelet endothelial cell adhesion molecule-1 (PECAM-1) and observed an increase in PECAM-1 expression within the expanded cambium and in cellularized pockets within the cortical bone (Fig. 6E).

### Elements lacking a periosteum do not grow outward

Our data suggests that the abnormal woven bone observed in PCC mice is driven by over-proliferation of osteoprogenitors in the periosteum while dynamic histomorphometry also revealed an increase in endosteal mineralization. We therefore predicted that limb skeletal elements that lack a periosteum would increase bone mass internally but not from the outer surface. To test this, we examined the morphology of two distinct *Prx1* lineage-derived (GFP+), skeletal elements in PCC and control mice without a periosteum: the patella in the hindlimb and carpals in the forelimb (Fig. 7A, C, G, E, S5). The patella is the largest and most accessible sesamoid element and is surrounded by fibrocartilaginous attachments to tendons and ligaments and by articular cartilage. Carpals are small bones of the wrist that are surrounded entirely by articular cartilage.

**Figure 7.**
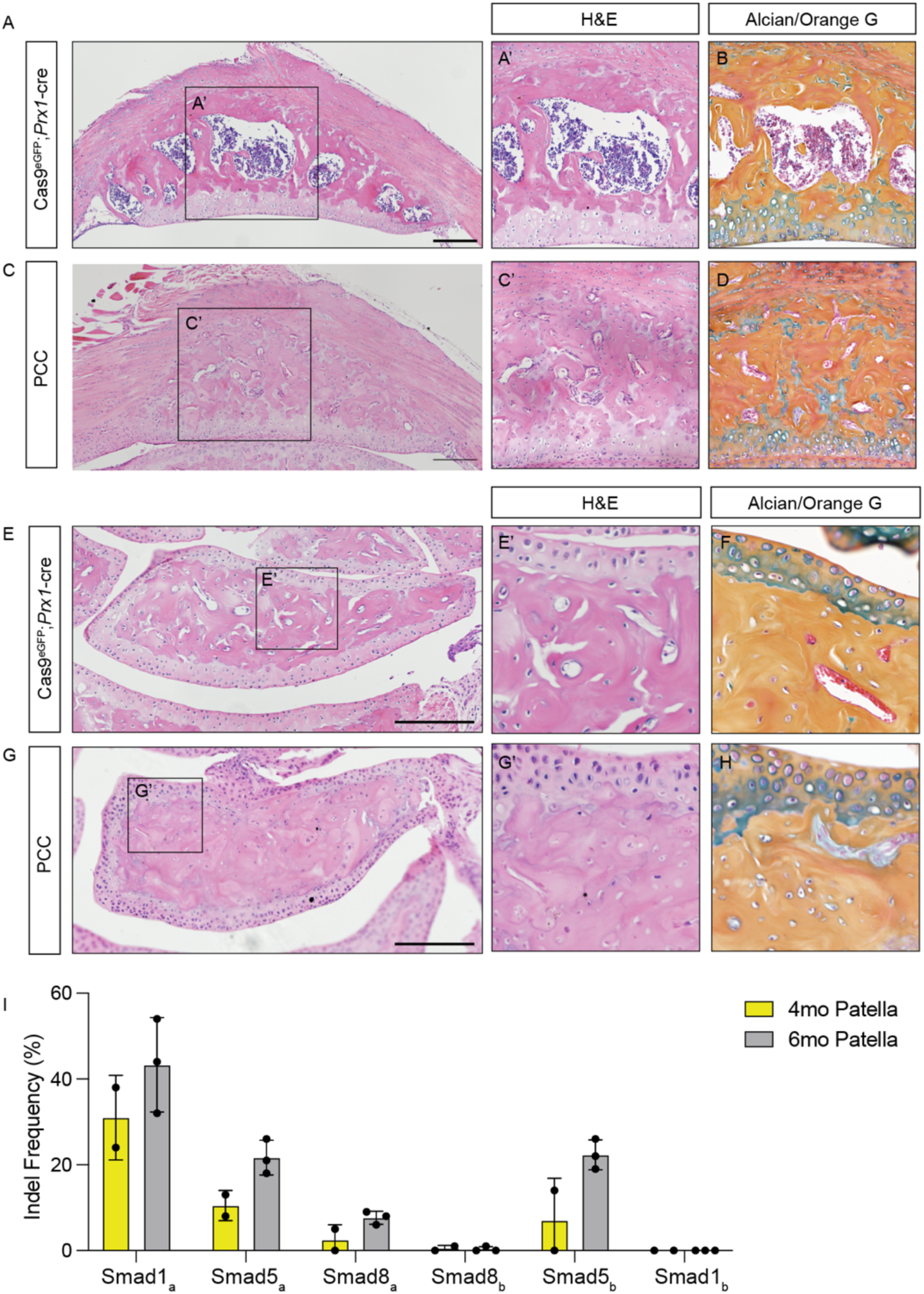
Patella and carpals, which lack a periosteum, do not exhibit surface bone overgrowth. **(A)** H&E staining of a patella section from four-month-old Cas9^eGFP^;*Prx1*-cre mouse and **(C)** four-month-old PCC mouse. Black boxes show higher magnification of marrow space **(A’)** and **(C’)**. **(B)** and **(D)** Alcian/Orange G stain of serial patella sections. **(E)** H&E staining of carpal sections from four-month-old Cas9^eGFP^;*Prx1*-cre mice and **(G)** four-month-old PCC mice. Black boxes show higher magnification of the inner-carpal space **(E’)** and **(G’)**. **(F)** and **(H)** Alcian/Orange G stain of serial carpal sections. (n=6 for Cas9^eGFP^;*Prx1-*cre and n=6 for PCC). **(I)** Indel analysis on patella tissue from four- and six-month-old PCC animals. Scale bars in A, C = 200 μm. Scale bars in E, G = 100 μm.

H&E and Alcian/Orange G staining of patella and carpal sections confirmed the absence of surface woven bone in these elements without a periosteum, and the marrow cavity of the patella is indeed filled with bone in *PCC* mice (Fig. 7C’, D). Similarly, although the carpals of control animals do not have a large marrow cavity like the patella, smaller vascular spaces are filled with bone in PCC mice at four months (Fig. 7G’, H).

To determine the degree to which non-periosteal elements accumulate mutations in *Smad/1/5/8,* we performed indel analysis on patella of four- and six-month mice (Fig. 7I). At four months, indel frequencies in *Smad1* and *Smad5* were 31% and 21.5%, respectively, while frequencies at *Smad8* were ∼5%. By six months this increased to 43.3%, 43.9%, and 8% for *Smad1*, *5*, and *8*. Together these data support the hypothesis that the surface woven bone expansion of long bones originates in the periosteum and that BMP/GDF signaling is also required to limit endosteal osteogenesis.

## Discussion

Here, we found an unexpected role for *Smad1/5/8* transcription factors to restrict hyperostosis in the limb skeleton. These findings were a serendipitous outcome of the unintended gradual accumulation of indel mutations postnatally, induced by conditional Cas9 expression and an array of gRNAs targeting each *Smad*.

Compared to zygotic loss-of-function, conditional approaches are mosaic whether by cre/lox recombination or by Cas9 mutagenesis. Despite this limitation, reproducible phenotypes can be highly informative, and replicability is a distinct advantage of the Cas9 mutagenesis approach over traditional cre/lox systems. We obtained over 160 PCC mice hemizygous for all three transgenes (gRNA array, conditional Cas9^eGFP^, and cre) compared to just one reported *Smad1/5/8* conditional knockout that combined six floxed/null alleles and cre^3^.

Indeed, hyperostosis is highly reproducible and penetrant along the lengths of all four limbs in all PCC mice, despite the expected mosaicism of conditionally induced Cas9 mutagenesis, and it requires all three transgenes. Reproducibility is especially important, because we found it challenging to detect a significant reduction of SMAD protein expression due to high variance in Western blot samples using a phospho-Smad1/5/8 antibody. This may be caused by the mosaicism of conditional mutagenesis and/or differences in the cellularity of bone in PCC compared to control animals, which might affect the amount of cellular versus extracellular protein. However, RNA-Seq at the onset of hyperostosis at one month showed *Smad1* and *Smad8* mRNA expression is lower, likely due to nonsense mediated decay^41^, and GO pathway analysis suggests TGF-β and BMP signaling is lower than controls. Furthermore, the worsening phenotype over months coincides with increasing indel frequencies at each *Smad* target site, and we detected no indel mutations at any of the most likely off-target gRNA binding sites in PCC animals. Together, these data suggest that the observed phenotype is indeed caused by mutations in *Smad1/5/8* with the caveat that we cannot at this time determine how many or which of the six *Smad* target sites are mutated at cellular resolution *in situ*.

We suspect that the gradual accumulation of indels in PCC mice is due to low gRNA expression from a single genome-encoded transgene. Other PTG systems were tested by plasmid transfection or injection, which floods the cell with transgene copies^10,14,28^. By contrast, rate-limiting gRNA expression from a single-copy transgene might require time to overcome the slow kinetics of assembling ribonucleoprotein particles^42^. Additionally or alternatively, multiple gRNAs competing to bind Cas9 could result in initially low indel frequencies^10,28^. We also note that the last gRNA in the PTG array, Smad1_b_, never produced indels at its target site despite cutting DNA when tested individually *in vitro*. Further optimization of a multiplex gRNA array strategy may overcome these limitations to boost mutagenesis efficiency.

Rather than targeting non-essential genes as proof of feasibility, such as the pigmentation pathway, we chose to test a genome-encoded PCC system in mice that targets genes in a pathway relevant to our broader research interests. However, canonical BMP signaling is required for cell survival and proliferation in the early limb bud^43,44^. Thus, in an animal that gradually accumulates mosaic mutations, cells with *Smad1/5/8* mutations could be outcompeted by wild-type cells during earlier limb and cartilage development. If so, such competition might further delay or preclude the accumulation of mutations. A requirement for BMP/GDF signaling to later repress periosteal osteoprogenitor proliferation, as we observe here, might then give mutant cells a competitive advantage and allow the later phenotype to emerge.

Regardless of these technical limitations, the progressively worsening hyperostosis in PCC mice is consistent with and provides some clarity to previous BMP/GDF pathway loss-of-function phenotypes. *Bmpr1a* knockouts in mature osteocytes using *Dmp1-*cre and *Sp7-*cre caused an increase in bone mass and increased the rate of bone formation in mice at P33 and P60^45,46^. Knockout of *Bmpr2* with *Prx1-*cre increased trabecular and cortical bone, bone mineral density, and osteoblast activity at nine and 15 weeks^47^. Lastly, *Bmp3* null mice exhibit a striking increase in bone volume and trabecular and cortical thickness at eight and 16 weeks^48^. However, increased bone mass in these contexts is typically composed of normal bone - cancellous in the trabecular region and lamellar in the cortical region.

These differ from most knockouts of BMP signaling components, which cause a reduction in bone mass presenting in younger animals if endochondral bone develops and knockout is not lethal^3,44,49^. However, two studies observed an age-dependent change in bone mass when the same knockout was examined in younger versus older animals. Young mice with ablated *Bmpr1a* in maturing osteoblasts have low bone mass due to reduced osteoblast activity, but this reversed at six months when aged mice had increased bone mass due to reduced resorption^50^. Similarly, a *Smad4* knockout using the same *Osteocalcin-*cre (*OC-*cre) initially displayed low trabecular bone mass until seven months, when bone volume increased^51^. Altogether, these data suggest that BMP signaling may switch from a positive regulator of osteogenesis during bone formation and growth to a negative regulator once the organ is established and must be maintained into adulthood. Our data support this hypothesis, because the later postnatal accumulation of *Smad1/5/8* mutations coincides with an increase in bone mass. Our dynamic histomorphometry also suggests that BMP signaling negatively regulates bone formation by different mechanisms in the periosteum and endosteum - by restricting osteoprogenitor proliferation and osteoblast activity, respectively.

To test the hypothesis that the periosteum is necessary for outward expansion of woven bone, we examined the patella and carpals by μCT and histology. These skeletal elements, which lack a periosteum and are instead surrounded by articular cartilage and/or fibrocartilaginous tissue, have a similar size and shape in PCC mice and controls. This suggests that the periosteum is indeed necessary for surface hyperostosis. By contrast, the marrow cavity of the patella and vascular spaces in carpals are nearly filled with bone by four months in PCC mice, consistent with the increased mineral apposition rate and bone forming rate on endocortical surfaces of long bones. Together, this supports a hypothesis that *Smad1/5/8* are required to limit proliferation of periosteal osteoprogenitors and to prevent woven bone expansion and also to restrict the mineralizing activity of endocortical osteoblasts. Follow up studies using cre-drivers to more narrowly induce *Smad1/5/8* conditional mutagenesis and better optimized PTG arrays will refine the timing and cell type contributions to these phenotypes.

## Acknowledgements

We thank Drs. Vicki Rosen, Aris Economides, Tamara Alliston, Edward Hsiao, Andrew McCulloch, Sam Ward, and all members of the Cooper laboratory and the Wu Tsai Human Performance Alliance at UCSD for thoughtful discussion. We also thank Dr. Robert Goulet at the University of Michigan for generating the Dragonfly (ORS) plugins used for bone morphometric analysis. μCT scanning with calibration standard was performed by Haley Martens and Ramon Nagesan at the University of Michigan Museum of Zoology, with the help of Dr. Talia Moore.

We thank the Biorepository and Tissue Technology Shared Resources Center at the Moores Cancer Center at UCSD for assistance with histology and the Molecular and Cellular Immunology Core Flow Cytometry Unit at the Veteran Affairs Hospital at UCSD. Fluorescent automated cell sorting was made possible with help from the San Diego Center for AIDS Research (SD CFAR), an NIH-funded program (P30 AI036214), which is supported by the following NIH Institutes and Centers: NIAID, NCI, NHLBI, NIA, NICHD, NIDA, NIDCR, NIDDK, NIMH, NIMHD, NINR, FIC, and OAR. This publication also includes data generated at the UC San Diego IGM Genomics Center utilizing an Illumina NovaSeq X Plus that was purchased with funding from a National Institutes of Health SIG grant (#S10 OD026929). This work was supported by the Pathways in Biological Sciences NIH training grant (T32 GM133351) awarded to R.L. and by the National Institutes of Health under award numbers R21 AR077799 and R01 GM148640.

## Methods

### Mice and PTG line generation

The Rosa26-LSL-Cas9^eGFP^ [Stock #026175; B6J.129(B6N)-*Gt(ROSA)26Sortm1 (CAG-cas9*,-EGFP)Fezh*/J] and *Prx1-*cre [Stock #005584; B6.Cg-Tg(Prrx1-cre)1Cjt/J] lines were purchased from the Jackson Laboratory (ME, USA). The *Smad1/5/8* PTG mouse line was generated at The Jackson Laboratory using Bxb1 Integrase Mediated Cassette Exchange (B-RMCE), as previously described^52^, by delivery of a donor plasmid (p5178) into heterozygous zygotes of the Rosa26 Dual Bxb1 Landing Pad mouse strain (JAX Stock #36152). Briefly, the donor plasmid was generated by commercial cloning (GenScript) of the synthesized PTG array (∼1.4kb) into the Donor Vector p5154 (ADDGENE #175390), positioned between the *SacI* and *AgeI* restriction enzyme sites. Sequences of each gRNA and the sequence of the complete PTG array can be found in Supplemental Tables 2 and 6, respectively. The sequence of the complete PTG array inserted into the Rosa26 locus can be found in Supplemental Table 7.

Genomic integration of the transgene was achieved by microinjection into fertilized oocytes of commercially prepared (TriLink BioTechnologies) Bxb1 Integrase mRNA (100 ng/µl), Donor Plasmid #5178 (30ng/µl), and RNasin® Ribonuclease Inhibitor (Promega, cat# N2515; at 0.2U/µl). Both Bxb1 mRNA and the Donor Plasmid were ultra-purified for microinjection using Phenol-Chloroform extraction and ethanol precipitation. All reagents for microinjection were prepared in nuclease-free TE (10mM Tris, 0.1mM EDTA, pH7.5; Integrated DNA Technologies, Inc.; cat# 11-05-01-05). Microinjected zygotes were transferred to pseudo-pregnant females and carried to term. From four transfers (total of 78 embryos), 23 mice were born (29% survival), and nine candidates were identified with the transgene insertion (39%). Only 1/23 (4%) of the mice born (1/9 of the transgenics) had evidence of an off-target insertion (i.e., random insertion transgenesis). Two male candidates (#5 and #9) were bred to wild-type C57BL6/J (JAX Stock #000664) to verify germline transmission and to establish N1 generation heterozygotes. Both founders successfully produced offspring and the transgene insertion was confirmed at only the Rosa26 site. Candidate #5 was also bred to females from B6.Cg-Tg(Prrx1-cre)1Cjt/J (JAX Stock #005584).

The Cas9^eGFP^ transgene and the PTG transgene are both inserted at the Rosa26 locus, therefore triple transgenic offspring could only carry one copy of each. Line-specific primers (Supplemental Table 3) were used to genotype each animal. Control mice used for this study were littermates of the genotypes PTG/Cas9^eGFP^ and *Cas9^eGFP^*;*Prx1-*cre. Male and female animals were used for all experiments, at least n=3 for each sex and genotype. All mice were bred on the C57BL6/J background, housed under specific pathogen-free (SPF) conditions, and provided with food and water *ad libitum*. All animal care and use protocols were approved by the Institutional Animal Care and Use Committee (IACUC) of the University of California San Diego.

### gRNA validation and indel efficiency analysis

gRNAs were designed using CHOPCHOP^53^ and synthesized as forward and reserve oligos by Integrated DNA Technologies (IA, USA). To validate and test the cutting efficiency of each gRNA, complementary oligos were ligated and cloned into px458 plasmid directly upstream of a gRNA scaffold at the *BbsI* site (Addgene plasmid #48138). Each plasmid was transfected into immortalized embryonic mouse fibroblasts (iMEFs) (ATCC, CRL-2907) using TransitX2 (Mirus, MIR 6004) in triplicate. iMEFs were incubated at 37°C for 72 hours post transfection in DMEM (ATCC, 30-2002) supplemented with 10% FBS (Omega Scientific, NC0471611) and 1% Pen/Strep (ThermoFisher, 15140122). Cells were then fluorescently sorted for GFP to isolate those expressing GFP from the plasmid. Flow cytometry was done through The San Diego Center for AIDS Research, Molecular and Cellular Immunology Core (P30 AI036214). Genomic DNA was extracted from sorted GFP+ cells and gRNA target sites were amplified with PCR using primers (Supplemental Tables 4 and 5) that were designed ∼100 bp up- and downstream of the target sites. PCR products were Sanger sequenced by GeneWiz (NJ, USA) and analyzed using Inference of CRISPR Edits (ICE) (EditCo). *In vivo* indel frequencies were tested similarly using bone tissue from the distal radius/ulna or patella. Tissue was collected from sacrificed animals at various time points, cleared of marrow and connective tissue, flash frozen, and ground with a mortar and pestle prior to genomic extraction. All subsequent steps were identical to those done for *in vitro* indel analysis. Off-target gRNA genomic locations were identified using three independent *in silico* prediction webtools, CHOPCHOP, Off-spoter, and Cas-OffFinder. Similar to target site indel analysis, primers were designed ∼100 bp up- and downstream of each predicted site, PCR amplified, Sanger sequenced, and analyzed with ICE. The same genomic DNA was used for both target and off-target site indel analysis.

### Western Blotting

The distal radius and ulna of PCC and control animals were dissected in ice cold PBS, carefully cleaned of muscle and connective tissue, and the marrow cavity was flushed with PBS. Samples were immediately flash frozen in liquid nitrogen and stored at −80 °C. For tissue disruption, bone samples were ground into a powder using a mortar and pestle over liquid nitrogen, then resuspended in RIPA Lysis and Extraction Buffer (Thermo, 89900) with Halt™ Protease and Phosphatase Inhibitor Cocktail (Thermo, 78440) used at 1X concentration and kept on ice during preparation. Bone fragments were sonicated for 10s at 50% power three times on a membrane dismembrator model 100 (Fisher Scientific) with a microtip probe then centrifuged for 10 min at 15,000 rpm at 4°C. Lysate protein concentrations were measured by BCA Protein Assay (23225, Thermo Scientific Pierce). Laemmli sample buffer with β-mercaptoethanol was then added to cell lysates and heated at 95°C for 5 min. Samples were then cooled to room temperature and centrifuged briefly. Lysates were resolved on Mini-PROTEAN TGX Stain-Free Precast Gels (Bio-Rad), followed by transfer to PVDF membranes (1620177, BioRad) using Bjerrum semi-dry transfer buffer (48 mM Tris Base, 39 mM Glycine-free acid, 0.0375% SDS, 20% MeOH, pH 9.2) and a semi-dry transfer apparatus (Bio-Rad Turbo Transfer) for 30 min at 25V. Immunoblots were blocked with 5% blotting grade nonfat dry milk (APEX Bioresearch) in TBST for 1 hour. Primary antibodies were diluted in 5% BSA and rocked overnight. The following antibodies were used: phospho-Smad1/5/8 (Cell Signaling, 13820), GFP (Thermo, A10262), B-actin (Cell Signaling, 4967), anti-rabbit IgG HRP (Cell Signaling, 7074), and anti-chicken IgG HRP (Thermo, A16057). Immunoblots were developed using Clarity Western ECL Substrate (BioRad, 1705061) and imaged on a Bio-Rad Chemi-Doc XRS+ system. All blots were processed using Bio-Rad Image Lab software (RRID:SCR_014210), with final images prepared in Adobe Illustrator.

### RNA-Sequencing and Analysis

The distal radius and ulna of PCC and control animals were dissected in ice cold PBS, carefully cleaned of muscle and connective tissue, and the marrow cavity was flushed with PBS. Samples were immediately placed in RNAlater (Invitrogen) and equilibrated overnight at 4 °C. Samples were removed from RNAlater and snap frozen in liquid nitrogen the next day and stored at −80 °C or immediately processed for RNA extraction.

For tissue disruption, bone samples were ground into a powder using a mortar and pestle over liquid nitrogen, then homogenized using the Qiagen QIAshredder kit per the manufacturer’s protocol. RNA was extracted from the tissue homogenate using the Qiagen RNeasy Micro kit following the manufacturer’s protocol, including the on-column DNA digest step. RNA concentrations were measured via Nanodrop before submission to the UCSD IGM core for tape station measurements. Samples with RIN scores >8 were used for library preparation. UCSD IGM used Illumina mRNA Stranded Library kits to prepare libraries. Samples were run on PE100 lanes to get 25 million reads per sample on an Illumina NovaSeq 6000 platform.

Adapter trimming and read quality control analysis were performed using Trim Galore, a wrapper around Cutadapt^54^ and FastQC^55^. Trimmed reads were mapped to the mouse genome (mm10) using STAR^56^, and read counts per gene were acquired by “–quantMode GeneCounts” option. STAR aligned gene counts were used to perform differential expression analysis by DESeq2^57^. Of four PCC and three PTG/Cas9 samples collected for the analysis, two PCC and one PTG/Cas9 were collected in batch A, and the others were collected in batch B. Therefore, we used ComBat-seq^58^ to account for potential batch effects prior to differential expression analysis. GO term enrichment analysis was performed using Enrichr^59^.

### μCT scanning and analysis

Forelimbs and hindlimbs of PCC mice and both control genotypes, PTG;Cas9^eGFP^ and Cas9^eGFP^;*Prx1*-cre, were collected in 70% ethanol and stored at 4°C until μCT scanning. Scanning was done at the University of Michigan Museum of Zoology µCT Scanning Laboratory (MI, USA). Forelimbs and hindlimbs were scanned using Nikon XT H 225ST µCT Scanner. The following settings were applied: 7 or 10 µm voxel Source: 225kV, Beam Energy:90kV, Beam Current: 200 µA, Exposure: 0.354 s, Filter: None. After scanning, 3D images were reconstructed using Nikon CT Pro3D Software (Version XT 5.4). A hydroxyapatite standard (HA=1.69 g/cm^3^) was included with each limb scan to calculate BMD. 2D and 3D analyses were done in Dragonfly (ORS). Trabecular analysis was done on the first 40 slices from just underneath the chondro-osseus junction of the radius. Cortical analysis was done on 40 slices 5 mm from distal tip of the radius. The bone morphometric parameters BV/TV, Tb.Sp, Tb.Th, Tb.N, Tb.BMD, Ct.BMD, Ct.BV, and Ct.Th were quantified using plugins written by Dr. Robert Goulet at University of Michigan. Nomenclature and abbreviations of µCT parameters were followed according to guidelines recommended by American Society for Bone and Mineral Research^60^.

### Bone histology

For histologic evaluation of the bone, forelimbs and hindlimbs were dissected, carefully cleaned of muscle tissue, fixed in 4% PFA overnight, and decalcified for 14 days in 20% EDTA pH 8.0 with solution changes every two days. Limbs were dehydrated through an ethanol gradient, cleared in xylene, and embedded in paraffin. Radii, carpals, and patella were longitudinally sectioned at 5 μm with a microtome (Leica RM 2165). Hematoxylin and eosin (H&E), tartrate-resistant acid phosphatase (TRAP), and Alcian/Orange G stains were done according to protocols from the University of Rochester Medical Center. Three sections per individual for each stain were imaged on an Olympus BX61 upright compound microscope. Growth plate heights and heights of individual growth plate zones were measured using the InteredgeDistance macro (Santosh Patnaik) in FIJI^61^. Osteoclast number was calculated using TrapHisto, an open-source image analysis software based in FIJI developed by Dr. Rob J. van’t Hof^62^. Results were visualized in Graphpad Prism.

### Skeletal preparations

Skeletal staining protocol was performed as previously described in^63^. Briefly, newborn (P0) animals were euthanized, and forelimbs and hindlimbs were dissected, skinned, and fixed in 4% PFA overnight, then moved to PBS. Samples were stained over two nights in cartilage staining solution (75% ethanol, 20% acetic acid, and 0.05% Alcian blue 8GX (Sigma-Aldrich, A3157)), rinsed overnight in 95% ethanol, cleared overnight in 0.8% KOH, and stained overnight in bone staining solution (0.005% Alizarin Red-S (Sigma-Aldrich, A5533) in 1% KOH). After staining, samples were further cleared in 20% glycerol in 1% KOH until digits were free of soft tissue and long-bone morphology was visible. Samples were further processed through a graded series of 50% and 80% glycerol in 1% KOH and then into 100% glycerol for imaging and storage. All steps of the skeletal staining procedure were performed with gentle rocking at room temperature.

### In vivo dynamic histomorphometry

P37 mice received an intraperitoneal injection of calcein (20 mg/kg, Sigma, C0875) dissolved in 1x PBS followed four days later by an intraperitoneal injection of alizarin (40 mg/kg, Sigma, A5533) dissolved in 1x PBS. Mice were sacrificed 1 day later; injections and collections were timed such that mice were six weeks old on the day of collection. Forelimbs were dissected and fixed in 4% PFA overnight, taken through a sucrose gradient, and embedded in Optimal Cutting Temperature (OCT) media. Samples were flash frozen in OCT media in block molds. 6 μm thick longitudinal serial sections were taken of the radius using a Leica CM 1950 cryostat. Slides were rinsed of OCT using 1x PBS for 5 minutes and incubated in DAPI for 2 minutes then mounted in Fluoromount-G Mounting Media. Three sections per individual were imaged using an inverted Olympus Fluoview 3000 laser scanning confocal microscope at 20x magnification. All images were taken at the mid-diaphysis, visualized in FIJI as maximum intensity projections, and cropped to the same size regions of interest (ROIs) for analysis in Osteomeasure Version 3.3 (Osteometrics). The following parameters were measured: bone surface length (BS); single label perimeter (sL.Pm); double-label perimeter (dL.Pm); and double label width (dL.Ith). From primary data, we derived the mineralizing surface: MS/BS = (1/2sL.Pm + dL.Pm)/B.Pm X 100%; mineral apposition rate: MAR = dL.Ith/4 days in μm/d; and bone formation rate: BFR/BS = MAR X MS/BS; μm^3^/μm^2^ per day.

### Immunofluorescence

Animals were sacrificed at specified timepoints and forelimbs and hindlimbs were dissected, carefully cleaned of muscle tissue, fixed in 4% PFA overnight, and decalcified for 14 days in 20% EDTA, pH 8.0 with solution changes every other day. After which, limbs were taken through a sucrose gradient, embedded in OCT, and flash frozen in block molds. Blocks were stored at -80°C until cryo-sectioned. Frozen blocks were cryo-tape sectioned at 10 μm thickness longitudinally using Leica tape sectioning solution and CryoJane materials.

For immunofluorescence, slides were washed for 5 min in 1x PBS and subject to antigen retrieval either by incubation in Proteinase K (5 μg/mL) for 10 min or Bioworld Epitope Unmasking Buffer (citric acid based) per the manufacturer’s protocol, depending on the primary antibody. After antigen retrieval, slides were post-fixed with 4% PFA followed by three washes in 1x PBS. Slides were then blocked in a solution of 5% heat inactivated goat serum, 3% BSA, 0.1% TritonX-100, 0.02% SDS in PBS. Slides were incubated in the appropriate primary antibody dilution in block overnight at 4°C. On the second day, slides were washed three times for 10 min in PBST (1x PBS + 0.1% TritonX-100) and incubated at room temperature in secondary antibodies and 1 µg/ml DAPI for 1 hr. Slides were then washed three times for 10 min in PBST and mounted in Fluoromount-G Mounting Media. Four sections per individual were imaged on an inverted Olympus Fluoview 3000 laser scanning confocal microscope. For each section the distal metaphysis and the mid-diaphysis of the radius/ulna were imaged. For all fluorescence-based imaging on the laser scanning confocal microscope, laser detection thresholds were set against a no-primary antibody control. For each image, 12 z-stacks were captured.

The following primary antibodies and dilutions were used for immunofluorescence of tissue sections: SCA-1(Invitrogen, 14-5981-82, 1: 250); POSTN (Abcam, ab14041, 1: 100) [citric acid]; PECAM (Invitrogen, 14-0311-82, 1: 250); GFP (Invitrogen, A10262, 1: 500); RUNX22 (Abcam, ab76956, 1:500); COL1A1 (Developmental Studies Hybridoma Bank, Sp1.D8, 1:20) [citric acid]; Ki67 rat (Invitrogen, 14-5698-82, 1: 150); Ki67 rabbit (Abcam, ab15580, 1:200); PRX1 (Invitrogen, PA5-106700, 1: 100). The following secondary antibodies and dilutions were used for immunofluorescence, all were obtained from Invitrogen: SCA1 – goat ranti-rat IgG 594 (1:250); POSTN - goat anti-rabbit IgG 647 (1:500); PECAM – donkey anti-goat 594 (1:500); GFP - goat anti-chicken IgG 488 (1:500); RUNX2 – goat anti-rabbit IgG 647 (1:500); COL1A1 – goat anti-mouse IgG1 (1:250); Ki67 – goat anti-rat IgG 594 (1:1000); PRX1 – goat anti-rabbit IgG 647 (1:800).

For TUNEL staining, slides were generated with the same protocol as for immunofluorescence and stored at -80°C until staining. Slides were rinsed in 1x PBS for five minutes and placed immediately into the TUNEL reaction mixture following manufacturer’s instructions (Roche In Situ Cell Death Detection Kit, TM-Red) for 60 min at 37°C, rinsed three times in 1x PBS, and mounted in Fluoromount-G Mounting Media.

Raw images were opened in FIJI and prepared for bulk processing in Adobe Photoshop by separating channels and setting appropriate LUTs. For cell number quantifications in FIJI, images were cropped to only the regions of interest (ROI) (distal metaphysis or mid-diaphysis) using only the DAPI channel. ROI images were thresholded in each channel to create a mask. Watershed function was applied to separate adjacent cells and ‘Analyze Particles’ function was used to count each size-restricted shape. The cell size for all analyses was set to 5 μm. To calculate the proliferative index, the ratio of Ki67 positive to total number of cells per ROI was averaged across four sections from each individual at both the distal metaphysis and mid-diaphysis.

For co-localization analysis, ROI images were once again thresholded in each channel to create a mask, and the watershed function was applied to separate adjacent cells. The image calculator function was used with the ‘AND’ command to identify which cells expressed both Ki67 and COL1A1, for example. ‘Analyze Particles’ function was used to count each size-restricted, double-positive cell. Results were visualized in Graphpad Prism.

For each representative image, channels were organized as layers in Photoshop, grouped by fluorophore (ie. DAPI, COL1A1, etc.), and set as ‘screens’ against a black background. The levels tools was used to adjust image brightness and contrast equally across each antibody stain. After batch processing, images were moved to Adobe Illustrator for figure construction.

### Statistical analysis

Results were expressed as means ± SD. Comparisons between control and experimental animals were analyzed using two-tailed unpaired Student’s t-test and a *p* value of less than 0.05 was considered to indicate statistical significance. Three males and three females were used for each experiment per genotype for a total of six biological replicates (unless otherwise stated).

Quantifications of cell number and double positive cells were done on both the distal metaphysis and mid diaphysis, but only representative images and values for the mid-diaphysis are reported (unless otherwise stated). Briefly, for each animal, four serial sections of the same tissue were imaged, analyzed for the respective quantification, and averaged to yield one representative value per individual.

