## Supplemental Figures 1-5 and Supplemental Tables 1-7 for "CRISPR-mediated conditional mutagenesis of *Smad1/5/8* reveals BMP/GDF signaling restricts postnatal bone overgrowth"

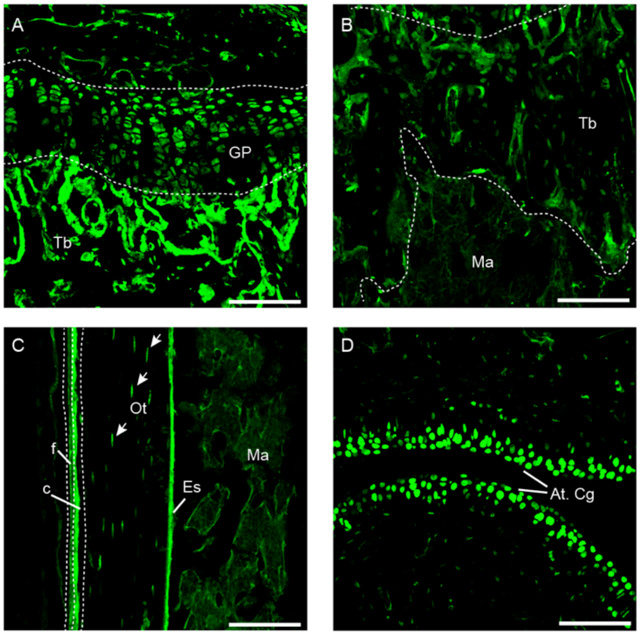


**Supplemental Figure 1 - *Prx1*-cre lineage trace revealing GFP+ cells as a proxy for Cas9 expression.**

Representative images from two-month Cas9$^{eGFP}$;*Prx1*-cre animals showing GFP expression throughout bone tissue. GFP was used as a proxy for Cas9 expression from the Rosa26-LoxSTOPLox-Cas9$^{eGFP}$ (Cas9$^{eGFP}$) transgene recombined by *Prx1*-cre. GFP expression was observed in growth plate chondrocytes (A), trabecular bone (A, B), the marrow cavity (B,C), cambium and fibrous layers of the periosteum (C), endosteum (C), embedded osteocytes (C, white arrowheads), and articular cartilage (D). GP: growth plate. Tb: trabecular bone. Ma: marrow cavity. c: cambium. f: fibrous. Ot: Osteocyte. Es: Endosteum. At. Cg: articular cartilage. Scale bar: 100 μm

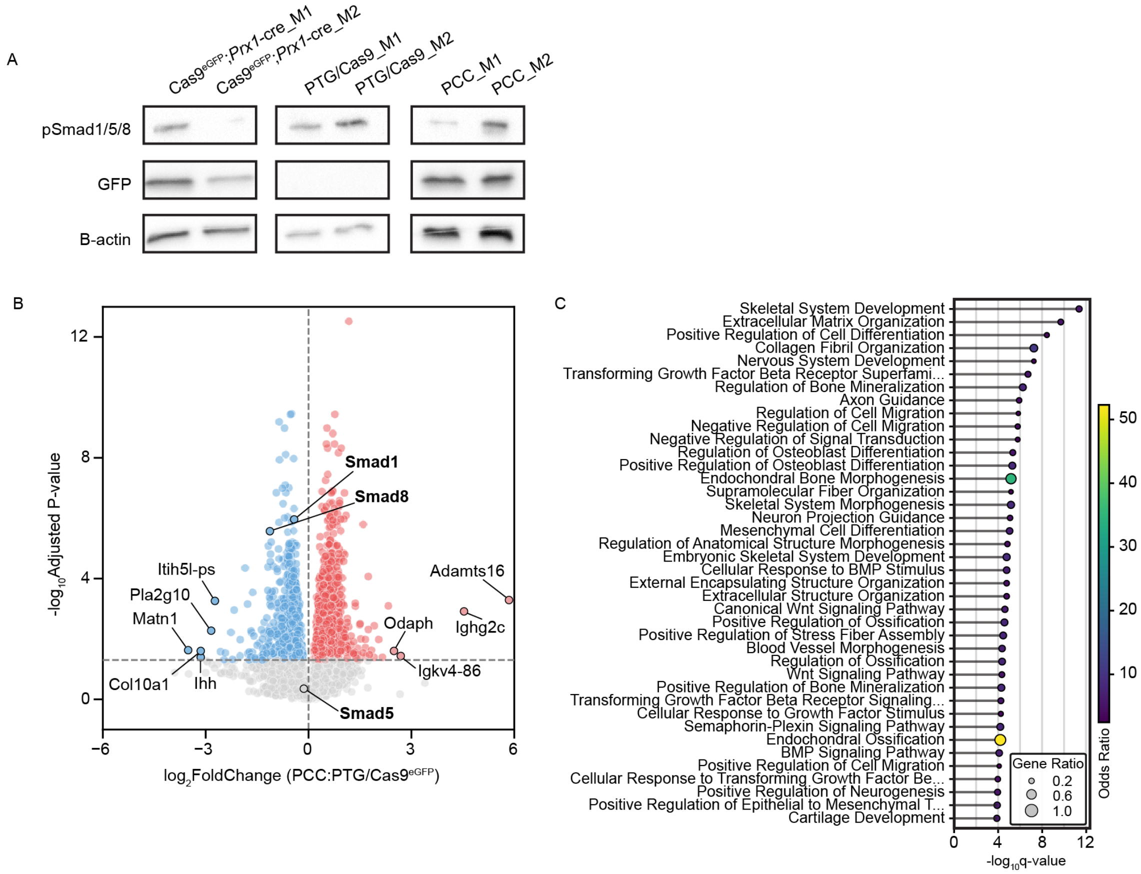


**Supplemental Figure 2 - Validation of reduced Smad1/5/8 activity**

**(A)** Variance in phospho-Smad1/5/8 Western blot results is difficult to quantify. **(B)** Bulk RNA-Seq of PCC (n=4) and PTG/Cas9$^{eGFP}$ (n=3) distal radius shows significantly reduced mRNA expression of *Smad1* and *Smad8* in PCC bone compared to control. **(C)** Enrichment analysis of GO biological process terms shows that TGF-β and BMP signaling pathways are among the most significantly enriched biological processes among all downregulated genes.

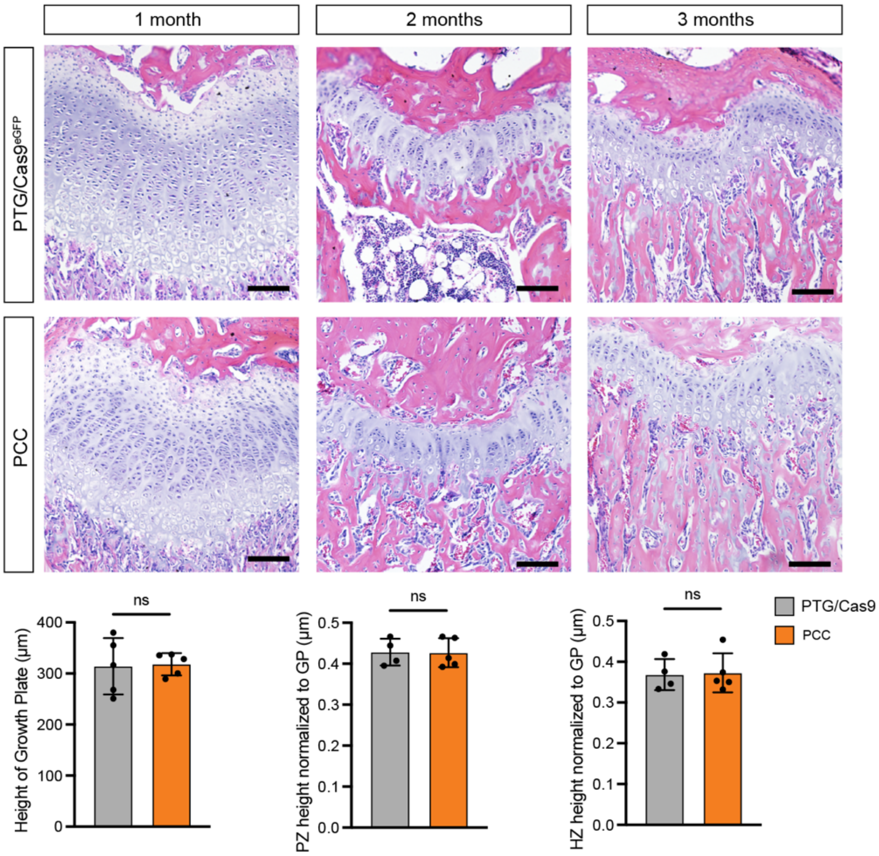


**Supplemental Figure 3 - Growth plate morphology is unchanged in PCC mice.**

H&E stain of growth plates from one-, two-, and three-month-old animals. Morphology, growth plate height, and height of individual zones (normalized to growth plate) are not significantly different between experimental and control animals. Quantifications are at one month. n=5 each for one-month PTG/Cas9$^{eGFP}$ and PCC. n=5 each for two-month PTG/Cas9$^{eGFP}$ and PCC. n=5 each for three-month PTG/Cas9$^{eGFP}$ and PCC*.* Scale bar = 100 μm.

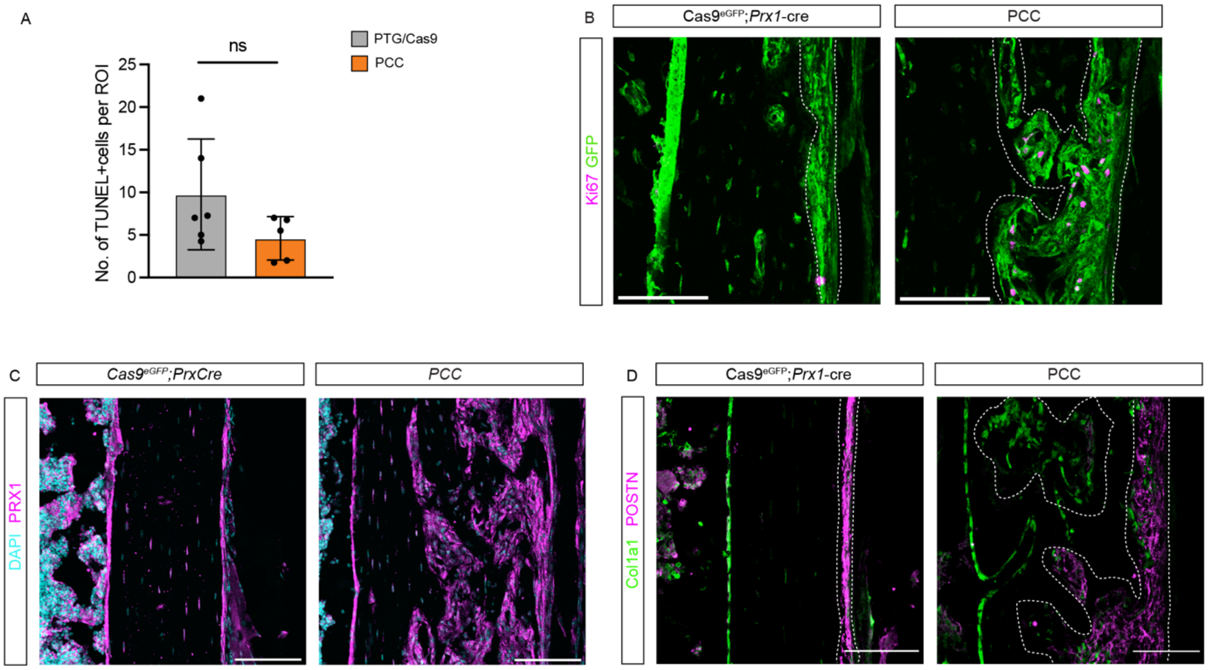


**Supplemental Figure 4 - Cellular characterization of expanded tissue in PCC mice.**

**(A)** Quantification of TUNEL positive cells from two-month-old mid-diaphysis of control and PCC mice. **(B)** Ki67 (magenta) and GFP (green) double immunofluorescence stain on two-month-old distal metaphyses. Periosteum is thicker in controls at the distal metaphysis compared to the mid-diaphysis. Ki67+ cells are present only throughout the expanded GFP+ periosteum. **(C)** PRX1 (magenta) immunofluorescence at the mid-diaphysis in two-month-old mice shows PRX1 expression throughout all the expanded tissue in PCC mice. **(D)** COL1A1 (green) and Periostin (POSTN, magenta) double immunofluorescence stain at the mid-diaphysis in two-month-old mice. The representative image shows expanded POSTN+ cambium cells that become embedded within cortical bone as pockets containing cells that differentiate into COL1A1+ osteoblasts and lose periosteal (POSTN+) identity. Data represent mean +/- SD. * denotes Welch's t-test <0.05. ** denotes Welch's t-test <0.005. *** denotes Welch's t-test <0.0005. ns = not significant. Scale bar = 100 µm.

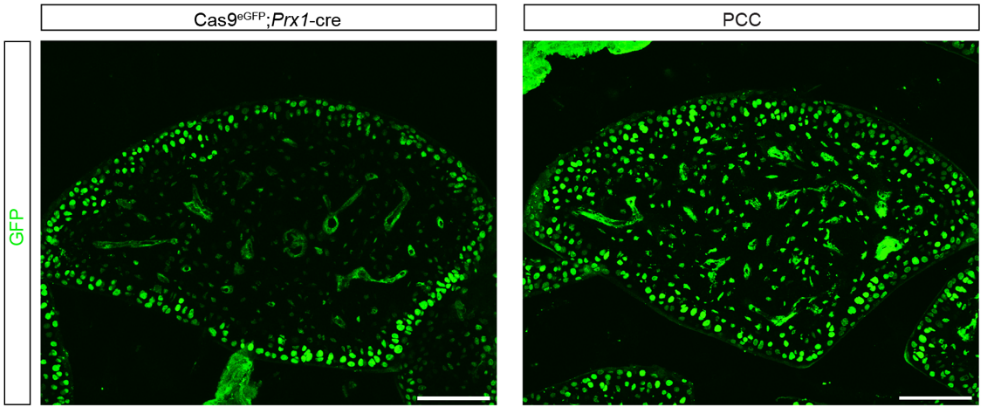


**Supplemental Figure 5 – GFP expression, a marker of the *Prx1*-lineage, in carpal elements.**

Carpal sections stained for GFP from two-month-old control and PCC animals. Scale bar = 100 μm.

**Supplemental Table 1 - Off-target ICE Analysis**

| Animal ID | Smad1(a) Ch11 | Smad1(a) Ch15 | Smad5(a) Ch5 |
|---|---|---|---|
| **L-272 (PCC)** | 0 | 0 | 0 |
| **L-538 (PCC)** | 0 | 0 | 0 |
| **L-539 (PCC)** | 0 | 0 | 0 |
| **L-287 (PCC)** | 0 | 0 | 0 |
| **L-264 (PCC)** | 0 | 0 | 0 |
| **L-492 (PCC)** | 0 | 0 | 0 |
| **L-454 (PTG/Cas9)** | 0 | 0 | 0 |
| **L-544 (PTG/Cas9)** | 0 | 0 | 0 |
| **L-279 (PTG/Cas9)** | 0 | 0 | 0 |
| **L-288 (PTG/Cas9)** | 0 | 0 | 0 |
| **L-618 (PTG/Cas9)** | 0 | 0 | 0 |
| **L-149 (PCC)** | 0 | 0 | 0 |
| **L-159 (PCC)** | 0 | 0 | 0 |
| **L-160 (PCC)** | 0 | 0 | 0 |
| **L-150 (PTG/Cas9)** | 0 | 0 | 0 |
| **L-158 (PTG/Cas9)** | 0 | 0 | 0 |
| **L-161 (PTG/Cas9)** | 0 | 0 | 0 |
| **L-155 (C9;Cre)** | 0 | 0 | 0 |
| **L-146 (C9;Cre)** | 0 | 0 | 0 |
| **01_67_17 (PCC)** | 0 | 0 | 0 |

3-month animals in green
6-month animals in blue
18-month animals in yellow

**Supplemental Table 2 - gRNA sequences.**

| gRNA Name | Sequence (5' 3') |
|---|---|
| Smad1a | GGGAACTCACAGCATTCCAG |
| Smad5a | GACGTCCATCCAAGGACCTG |
| Smad8a | CGGGAACTCACAGCACTCCA |
| Smad8b | aCACCTGGAGGCGTCCATCG |
| Smad5b | AACTTGCAGACGTCCATCCA |
| Smad1b | GCCATCCAGGGAGCGAGGAA |

**Supplemental Table 3 - Genotyping Primers.**

| Oligo Name | Location | Sequence (5’ 3’) | PCR Settings |
|---|---|---|---|
| PTG_fwd | In 3’ end of PTG array | GGTGCACGGTTTACCACTGA | 58°C Anneal 15 sec<br>72°C Extension 2 min |
| PTG_Rev | In Right Homology arm of Rosa26 locus | CAGCCTCGATTTGTGGTGTA | |
| Cas9_Fwd | In LoxP site of $Cas9^{eGFP}$ transgene | TTAGGTCCCTCGACCTGCAG | 60°C Anneal 15 sec<br>72°C Extension 2 min |
| Cas9_Rev | In Cas9 | CTGTCGAACAGCAGGGCTC | |
| Prx1_Fwd | In Prx1 promoter | AGGGGGTCTGTAAAACGTCA | 60°C Anneal 15 sec<br>72°C Extension 2 min |
| Prx1_Rev | In cre gene | gtgaaacagcattgctgtcactt | |

**Supplemental Table 4 - Primers for Inference of CRISPR Edits.**

| Oligo Name | Sequence (5’ 3’) | PCR Settings |
|---|---|---|
| Smad1_M1_Fwd | ACAGGTGCTTGTTACAGCGA | 60°C Anneal 15 sec<br>72°C Extension 1 min 30 sec |
| Smad1_M1_Rev | GGGTGGAAACAGGGCGATGA | |
| Smad1_M1_FwdSeq | GGACATCTGAGCATGCCTGC | Used for sequencing |
| Smad1_M1_RevSeq | TGTGGCACAGTTGGAGCTAA | |
| Smad5_M1_Fwd | CGCCTCCACTTGGACTTTCT | 60°C Anneal 15 sec<br>72°C Extension 1 min |
| Smad5_M1_Rev | AGGCATGGGGTGTTCACAAA | |
| Smad5_M1_FwdSeq | GTGACTTTTCACCTGCGAGC | Used for sequencing |
| Smad5_M1_RevSeq | CAGCCTTTTCGCTAATAAGCC | |
| Smad8_M1_Fwd | TTGGAAGACGTCAGCCCATC | 60°C Anneal 15 sec<br>72°C Extension 1 min 30 sec |
| Smad8_M1_Rev | CCAACCTTGCAAGCCTTTCG | |
| Smad8_M1_FwdSeq | GAGCTCTGCCTCCTATGCAC | Used for sequencing |
| Smad8_M1_RevSeq | ACGGGCTCACACCCCGAATA | |

**Supplemental Table 5 - Primers for Inference of CRISPR Edits of Off-Target (OT) Sites.**

| **Oligo Name** | **Location** | **Sequence (5' 3')** |
|---|---|---|
| Smad1_Ch15OT_Fwd | Chromosome 15 | ATGTGGGCTGCATCTGGAC |
| Smad1_Ch15OT_Rev | Chromosome 15 | AGCTGGTGACGAGATCACTGG |
| Smad1_Ch15OT_Seq | Chromosome 15 | GGCCCAGCTGTGCTCACTAG |
| Smad1_Ch11OT_Fwd | Chromosome 11 | TTCCTGCCTCTCAGACCACAG |
| Smad1_Ch11OT_Rev | Chromosome 11 | TGGCAAGAGAATTACCCGGGG |
| Smad1_Ch11OT_Seq | Chromosome 11 | CGGCCAAGTGTCATCTGCTG |
| Smad5_Ch5OT_Fwd | Chromosome 5 | GACTTGCAGAGCTCACACTTAGGA |
| Smad5_Ch5OT_Rev | Chromosome 5 | CCTGCAGGAGATGCCTAGGG |
| Smad5_Ch5OT_Seq | Chromosome 5 | GTAGGAGGCAATCCTTGGCC |

**Supplemental Table 6 - Full sequence of PTG array.**

| | |
|---|---|
| **Full sequence of PTG array** | >GAGGGCCTATTTCCCATGATTCCTTCATATTTGCATATACGATACAAG<br>GCTGTTAGAGAGATAATTGGAATTAATTTGACTGTAAACACAAAGATAT<br>TAGTACAAAATACGTGACGTAGAAAGTAATAATTTCTTGGGTAGTTTGC<br>AGTTTTAAAATTATGTTTTAAAATGGACTATCATATGCTTACCGTAACTT<br>GAAAGTATTTCGATTTCTTGGCTTTATATATCTTGTGGAAAGGACGAAA<br>CACCAACAAAGCACCAGTGGTCTAGTGGTAGAATAGTACCCTGCCACG<br>GTACAGACCCGGGTTCGATTCCCGGCTGGTGCAGGGAACTCACAGCA<br>TTCCAGGTTTTAGAGCTAGAAATAGCAAGTTAAAATAAGGCTAGTCCGT<br>TATCAACTTGAAAAAGTGGCACCGAGTCGGTGCAACAAAGCACCAGTG<br>GTCTAGTGGTAGAATAGTACCCTGCCACGGTACAGACCCGGGTTCGAT<br>TCCCGGCTGGTGCAGACGTCCATCCAAGGACCTGGTTTTAGAGCTAGA<br>AATAGCAAGTTAAAATAAGGCTAGTCCGTTATCAACTTGAAAAAGTGGC<br>ACCGAGTCGGTGCAACAAAGCACCAGTGGTCTAGTGGTAGAATAGTAC<br>CCTGCCACGGTACAGACCCGGGTTCGATTCCCGGCTGGTGCACGGGA<br>ACTCACAGCACTCCAGTTTTAGAGCTAGAAATAGCAAGTTAAAATAAGG<br>CTAGTCCGTTATCAACTTGAAAAAGTGGCACCGAGTCGGTGCAACAAA<br>GCACCAGTGGTCTAGTGGTAGAATAGTACCCTGCCACGGTACAGACCC<br>GGGTTCGATTCCCGGCTGGTGCAaCACCTGGAGGCGTCCATCGGTTTT<br>AGAGCTAGAAATAGCAAGTTAAAATAAGGCTAGTCCGTTATCAACTTGA<br>AAAAGTGGCACCGAGTCGGTGCAACAAAGCACCAGTGGTCTAGTGGT<br>AGAATAGTACCCTGCCACGGTACAGACCCGGGTTCGATTCCCGGCTG<br>GTGCAAACTTGCAGACGTCCATCCAGTTTTAGAGCTAGAAATAGCAAG<br>TTAAAATAAGGCTAGTCCGTTATCAACTTGAAAAAGTGGCACCGAGTCG<br>GTGCAACAAAGCACCAGTGGTCTAGTGGTAGAATAGTACCCTGCCACG<br>GTACAGACCCGGGTTCGATTCCCGGCTGGTGCAGCCATCCAGGGAGC<br>GAGGAAGTTTTAGAGCTAGAAATAGCAAGTTAAAATAAGGCTAGTCCGT<br>TATCAACTTGAAAAAGTGGCACCGAGTCGGTGCTTTTT |

U6 promoter: in grey
Last 8bp of U6 promoter: white background
tRNA: in yellow
gRNA: in red
scaffold: in blue
Pol III Poly-T terminator: in purple

Note: Smad8b gRNA was incorrectly synthesized. The lowercase 'a' in Smad8b sequence is not supposed to be there, and there is an A missing at the end of the sequence.

**Supplemental Table 7 - Full Sequence of PTG Array insertion confirmed in Rosa26 locus.**

| | |
|---|---|
| **Full Sequence of PTG Array insertion confirmed in Rosa26 locus** | >GCGGCAGGCCCTCCGAGCGTGGTGGAGCCGTTCTGTGAGACAGCC<br>GGGTACGAGTCGTGACGCTGGAAGGGGCAAGCGGGTGGTGGGCAG<br>GAATGCGGTCCGCCCTGCAGCAACCGGAGGGGGAGGGAGAAGGGA<br>GCGGAAAAGTCTCCACCGGACGCGGCCATGGCTCGGGGGGGGGGG<br>GGCAGCGGAGGAGCGCTTCCGGCCGACGTCTCGTCGCTGATTGGCT<br>TCTTTTCCTCCCGCCGTGTGTGAAAACACAAATGGCGTGTTTTGGTTG<br>GCGTAAGGCGCCTGTCAGTTAACGGCAGCCGGAGTGCGCAGCCGCC<br>GGCAGCCTCGCTCTGCCCACTGGGTGGGGCGGGAGGTAGGTGGGGT<br>GAGGCGAGCTGGACGTGCGGGCGCGGTCGGCCTCTGGCGGGGCGG<br>GGGAGGGGAGGGAGGGTCAGCGAAAGTAGCTCGCGCGCGAGCGGC<br>CGCCCACCCTCCCCTTCCTCTGGGGGAGTCGTTTTACCCGCCGCCGG<br>CCGGGCCTCGTCGTCTGATTGGCTCTCGGGGCCCAGAAAACTGGCCC<br>TTGCCATTGGCTCGTGTTCGTGCAAGTTGAGTCCATCCGCCGGCCAG<br>CGGGGGCGGCGAGGAGGCGCTCCCAGGTTCCGGCCCTCCCCTCGG<br>CCCCGCGCCGCAGAGTCTGGCCGCGCGCCCCTGCGCAACGTGGCAG<br>GAAGCGCGCGCTGGGGGCGGGGACGGGCAGTAGGGCTGAGCGGCT<br>GCGGGGCGGGTGCAAGCACGTTTCCGACTTGAGTTGCCTCAAGAGG<br>GGCGTGCTGAGCCAGACCTCCATCGCGCACTCCGGGGAGTGGAGGG<br>AAGGAGCGAGGGCTCAGTTGGGCTGTTTTGGAGGCAGGAAGCACTTG<br>CTCTCCCAAAGTCGCTCTGAGTTGTTATCAGTAAGGGAGCTGCAGTGG<br>AGTAGGCGGGGAGAAGGCCGCACCCTTCTCCGGAGGGGGGAGGGG<br>AGTGTTGCAATACCTTTCTGGGAGTTCTCTGCTGCCTCCTGGCTTCTG<br>AGGACCGCCCTGGGCCTGGGAGAATCCCTTCCCCCTCTTCCCTCGTG<br>ATCTGCAACTCCAGTCTTTCTAGAACAAACCGGTTTGTCTGGTCAACC<br>ACCGCGGTCTCCGTCGTCAGGATCATCCAAGCTTAGATCTGAGTACTC<br>GAGCTCGAGGGCCTATTTCCCATGATTCCTTCATATTTGCATATACGAT<br>ACAAGGCTGTTAGAGAGATAATTGGAATTAATTTGACTGTAAACACAAA<br>GATATTAGTACAAAATACGTGACGTAGAAAGTAATAATTTCTTGGGTAG<br>TTTGCAGTTTTAAAATTATGTTTTAAAATGGACTATCATATGCTTACCGT<br>AACTTGAAAGTATTTCGATTTCTTGGCTTTATATATCTTGTGGAAAGGA<br>CGAAACACCAACAAAGCACCAGTGGTCTAGTGGTAGAATAGTACCCTG<br>CCACGGTACAGACCCGGGTTCGATTCCCGGCTGGTGCAGGGAACTCA<br>CAGCATTCCAGGTTTTAGAGCTAGAAATAGCAAGTTAAAATAAGGCTA<br>GTCCGTTATCAACTTGAAAAAGTGGCACCGAGTCGGTGCAACAAAGCA<br>CCAGTGGTCTAGTGGTAGAATAGTACCCTGCCACGGTACAGACCCGG<br>GTTCGATTCCCGGCTGGTGCAGACGTCCATCCAAGGACCTGGTTTTA<br>GAGCTAGAAATAGCAAGTTAAAATAAGGCTAGTCCGTTATCAACTTGA<br>AAAAGTGGCACCGAGTCGGTGCAACAAAGCACCAGTGGTCTAGTGGT<br>AGAATAGTACCCTGCCACGGTACAGACCCGGGTTCGATTCCCGGCTG<br>GTGCACGGGAACTCACAGCACTCCAGTTTTAGAGCTAGAAATAGCAAG<br>TTAAAATAAGGCTAGTCCGTTATCAACTTGAAAAAGTGGCACCGAGTC<br>GGTGCAACAAAGCAC |

**Supplemental Table 7 - Full Sequence of PTG Array insertion confirmed in Rosa26 locus (continued).**

| | |
|---|---|
| **Full Sequence of PTG Array insertion confirmed in Rosa26 locus (continued)** | CAGTGGTCTAGTGGTAGAATAGTACCCTGCCACGGTACAGACCCGGGTTC<br>GATTCCCGGCTGGTGCAaCACCTGGAGGCGTCCATCGGTTTTAGAGCTAG<br>AAATAGCAAGTTAAAATAAGGCTAGTCCGTTATCAACTTGAAAAAGTGGCA<br>CCGAGTCGGTGCAACAAAGCACCAGTGGTCTAGTGGTAGAATAGTACCCT<br>GCCACGGTACAGACCCGGGTTCGATTCCCGGCTGGTGCAAACTTGCAGA<br>CGTCCATCCAGTTTTAGAGCTAGAAATAGCAAGTTAAAATAAGGCTAGTCC<br>GTTATCAACTTGAAAAAGTGGCACCGAGTCGGTGCAACAAAGCACCAGTG<br>GTCTAGTGGTAGAATAGTACCCTGCCACGGTACAGACCCGGGTTCGATTC<br>CCGGCTGGTGCAGCCATCCAGGGAGCGAGGAAGTTTTAGAGCTAGAAAT<br>AGCAAGTTAAAATAAGGCTAGTCCGTTATCAACTTGAAAAAGTGGCACCGA<br>GTCGGTGCTTTTTACATCAGGTTGTTTTTCTGTTTTTACATCAGGTTGTTTT<br>TCTGTTTGGTTTTTTTTTTACACCACGTTTATACGCCGGTGCACGGTTTAC<br>CACTGAAAAACCGGTACTATgaattcgatatcGGATCCGATTTAAATTCGCGTAA<br>TCATGGGCCGGCTTGTCGACGACGGCGGACTCAGTGGTGTACGGTACAA<br>ACCAGGTAGATTAAAGACATGCTCACCCGAGTTTTATACTCTCCTGCTTGA<br>GATCCTTACTACAGTATGAAATTACAGTGTCGCGAGTTAGACTATGTAAGC<br>AGAATTTTAATCATTTTTAAAGAGCCCAGTACTTCATATCCATTTCTCCCGC<br>TCCTTCTGCAGCCTTATCAAAAGGTATTTTAGAACACTCATTTTAGCCCCAT<br>TTTCATTTATTATACTGGCTTATCCAACCCCTAGACAGAGCATTGGCATTTT<br>CCCTTTCCTGATCTTAGAAGTCTGATGACTCATGAAACCAGACAGATTAGT<br>TACATACACCACAAATCGAGGCTGTAGCTGGGGCCTCAACACTGCAGTTC<br>TTTTATAACTCCTTAGTACACTTTTTGTTGATCCTTTGCCTTGATCCTTAATT<br>TTCAGTGTCTATCACCTCTCCCGTCAGGTGGTGTTCCACATTTGGGCCTAT<br>TCTCAGTCCAGGGAGTTTTACAACAATAGATGTATTGAGAATCCAACCTAA<br>AGCTTAACTTTCCACTCCCATGAATGCCTCTCTCCTTTTTCTCCATTTATAA<br>ACTGAGCTATTAACCATTAATGGTTTCCAGGTGGATGTCTCCTCCCCCAAT<br>ATTACCTGATGTATCTTACATATTGCCAGGCTGATATTTTAAGACATTAAAA<br>GGTATATTTCATTATTGAGCCACATGGTATTGATTACTGCTTACTAAAATTT<br>TGTCATTGTACA |

Rosa26 Left homology arm in light grey
U6 Promoter in dark green
tRNA: in yellow
gRNA: in red
scaffold: in blue
Rosa26 Right homology arm in dark grey
